# Finite afterload during asymmetrically cycled preload promotes fetal ventricular growth, maturation, and contractile function while suppressing fibrotic remodeling

**DOI:** 10.64898/2026.06.17.733029

**Authors:** Mong Lung Steve Poon, Gaetano J. Scuderi, Ayaan Jamali, Alyson Dang, Jonathan T. Butcher

## Abstract

Myocardial development requires precise modulation of its growth and maturation by the mechanical loads within cardiac cycle, including preload and afterload. However, the mechanisms by which these natural cardiac loads interact to simultaneously govern growth and maturation of the developing myocardium in both cellular and mesoscale levels remain poorly understood. Here, we developed a naturally engineered fetal ventricular tissue (NFVT) platform that enables the application of afterload under dynamic preload using cyclic stretching with an asymmetrical duty cycle (asymmetrically cycled preload) to better replicate the natural cardiac loading in chick NFVT. Our results showed that low afterload (LA) enhanced NFVT contractile function with sustained tissue growth and improved cardiomyocyte maturation while suppressing fibrosis phenotypes. These effects were associated with reduced YAP1 and NOTCH activation in cardiomyocytes and enhanced tissue architecture. In contrast, high afterload (HA) induced contractile impairment with fibrotic remodeling through activation of cardiomyocyte PIEZO1/YAP1 signaling and sustained fibroblast entanglement.

**Teaser:** Low afterload under asymmetrically cycled preload promotes NFVTs contractile function with sustained tissue growth and cardiomyocyte maturation while suppressing fibrosis phenotypes through regulation of cellular mechanotransduction, including minimal PIEZO1 expression and inhibited YAP1 and NOTCH activation in cardiomyocytes, and improvement of collective cellular organization.

## Introduction

The heart drives blood circulation in the body from the very first heartbeat during embryogenesis, acting as a pump that sustains and adapts to varying mechanical loads throughout development. These natural cardiac loads mainly include stretching of the ventricle during chamber filling in diastole (Preload) and pressure the ventricle must overcome to contract for blood ejection in systole (afterload) within the cardiac cycle (1,2). In between each cycle, a short chamber relaxation period exists. This transient relaxation along with diastole maintains an asymmetric cardiac duty cycle, in which two-thirds of the cardiac cycle corresponds to diastole and the remaining one-third to systole (3).

During development, the natural cardiac loads undergo adaptive changes that progressively shape ventricular structure and function through modulation of myocardial growth and maturation (4). Ventricular chamber development begins with the trabeculation of a thin, slowly proliferating myocardium in the linear heart tube as cardiomyocytes protrude into the ventricular lumen to form a complex meshwork of trabeculae (5,6). At this stage, the myocardium is exposed to relatively lower mechanical loading due to the establishment of primitive unidirectional circulation that is characterized by lower heart rate, blood pressure, and volumetric workload (7–9). As development proceeds, the linear heart tube remodels into a chambered structure. Trabeculae then compact and integrate with the compact myocardium, leading to increased myocardial mass, while trabecular cardiomyocytes acquire more mature sarcomeric organization (10,11). Consequently, the myocardium thickens and ventricular contractility increases, contributing to progressively higher preload and afterload. Mechanical loads continue to rise postnatally as compaction advances and cardiomyocytes undergo substantial hypertrophic growth, increasing in both length and thickness (12,13). Despite knowledge on the morphogenic events and loading conditions outlining myocardial development, insight on how mechanical loading orchestrates the natural developmental paradigm, from cellular function and phenotypes, celcell signaling, to mesoscale cellular organization, in coordinating growth and maturation of the developing myocardium remains limit. This further restricts our ability to understand its disruption in congenital heart disease, such as left ventricle noncompaction and hypoplastic left heart syndrome.

Engineered heart tissues provide an intuitive in *vitro* platform to replicate the natural mechanical environment of the heart. With cardiac cells embedded in a hydrogel cast between two elastomeric posts, engineered heart tissues enable controlled modeling of preload and afterload (14–16). The distance between the posts establishes a baseline uniaxial tension that mimics preload and promotes anisotropic cell alignment, whereas the post geometry and mechanical properties determine the afterload against which the tissue contracts. To more faithfully replicate the dynamic loading conditions of myocardium throughout development, various bioreactor systems have been developed to actively modulate preload and afterload. Preload modulation is often achieved by adjusting tissue length with stepwise or cyclic stretching. However, most of these systems restrict contractile shortening during stretching due to the use of fixed anchorage, resulting in an effectively infinite afterload that severely induces pathological remodeling despite an enhanced tissue maturation (17–24). Furthermore, the use of stepwise or continuous cyclic stretching omits the duty cycle asymmetry during cardiac cycle in *vivo*, compromising their physiological relevance (15,21,25). Afterload modulation can be attained by either fabricating supporting posts with different stiffness or dynamically tunning post stiffness with mechanical brace or piezoelectric actuator (23,26–28). Nonetheless, the effect of distinct afterloads multiplexing with dynamic preload using cyclic stretching at an asymmetrical duty cycle (asymmetrically cycled preload) has not yet been systematically investigated. A platform that replicates such natural cardiac loading condition in *vitro* in the context of fetal myocardium can help better decode the obscured mechanisms underlying myocardial development without complicating by the effect of tissue pathogenesis. In this regard, human induced pluripotent stem cell-derived cardiomyocytes are increasingly used to develop various engineered heart tissue-based models and show great promise in developmental and disease modeling, drug screening, and cardiac regeneration. However, cardiac developmental models using these cardiomyocytes often fall short in recapitulating key aspects of myocardial development partly because of the lack in native multicellular complexity, maturation capacity, and developmental stage specificity(14,29,30).

In this study, we report the development of naturally engineered fetal ventricular tissue (NFVT) platform designed to better mimic the natural cardiac loading condition using distinct afterloads under asymmetrically cycled preload for the mechanical loading of chick NFVTs. The goal is to understand how natural cardiac loads alter myocardial growth, maturation, and pathological remodeling during development at both cellular and mesoscale levels. We examined tissue contractility, growth, structure organization, cardiomyocyte maturation, and pathological remodeling of NFVTs exposed to low afterload (LA) or high afterload (HA) under asymmetrically cycled preload. We then evaluated the potential role of cardiomyocyte PIEZO1, YAP1, and NOTCH mechanosignaling pathways underlying the effects of the two distinct loading regimens. Our results demonstrated that LA achieved superior contractile functions of NFVTs with sustained tissue growth, improved structure organization, enhanced cardiomyocyte maturation, and suppressed fibrosis phenotypes. These improvements were facilitated by the suppression of PIEZO1 and downregulation of YAP1 and NOTCH, in cardiomyocytes, consistent with the normal developmental program in chick myocardium. In contrast, NFVTs under HA showed fibrotic remodeling with structural and functional impairment, which was driven by cardiomyocyte PIEZO1/YAP1 coactivation.

## Results

### Development of naturally engineered ventricular tissue platform for the independent modulation of afterload and asymmetrically cycled preload

The naturally engineered ventricular tissue (NFVT) platform allowed the independent modulation of afterload and asymmetrically cycled preload applied on NFVT in a custom, 3D printed, high throughput, mechanical stimulation bioreactor. The bioreactor consisted of a linear actuator connected to a 3D printed tab grid system via a shaft coupling (Fig. 1A). This tab grid system was housed within a 3D printed well plate box placed on a 48-well plate and covered by a well plate lid secured with a lid attachment. The entire assembly was mounted on a 3D printed base holder with the linear actuator. Under this configuration, the tab grid system protruded individual tabs into each well of the 48-well plate containing NFVT cast between notches on the flexible and rigid posts of a PDMS post construct (Fig. 1, B-C). The flexible and rigid posts were spaced to yield NFVT ∼4 mm in length after compaction while the tab was positioned above NFVT. Each tab could supply asymmetrically cycled preload to NFVT by translating along with the linear actuator and repeatedly deflecting the flexible post above its notch, starting from a defined resting distance (d_rest_) between the tab and the flexible post and operating at specific strain percentage, linear speed, and thus duty cycle. The rigid post served as a reference point for cyclic stretching stabilized by a polylactic acid (PLA) strip. In contrast, afterload during tissue contraction was determined exclusively by the stiffness of the flexible post, which resisted tissue shortening as the tab retracted from flexible post and returned to its starting position in each loading cycle. Thus, asymmetrically cycled preload was governed by actuator-driven tab motion, whereas afterload was set by post stiffness, enabling independent modulation of the two parameters.

**Fig. 1.**
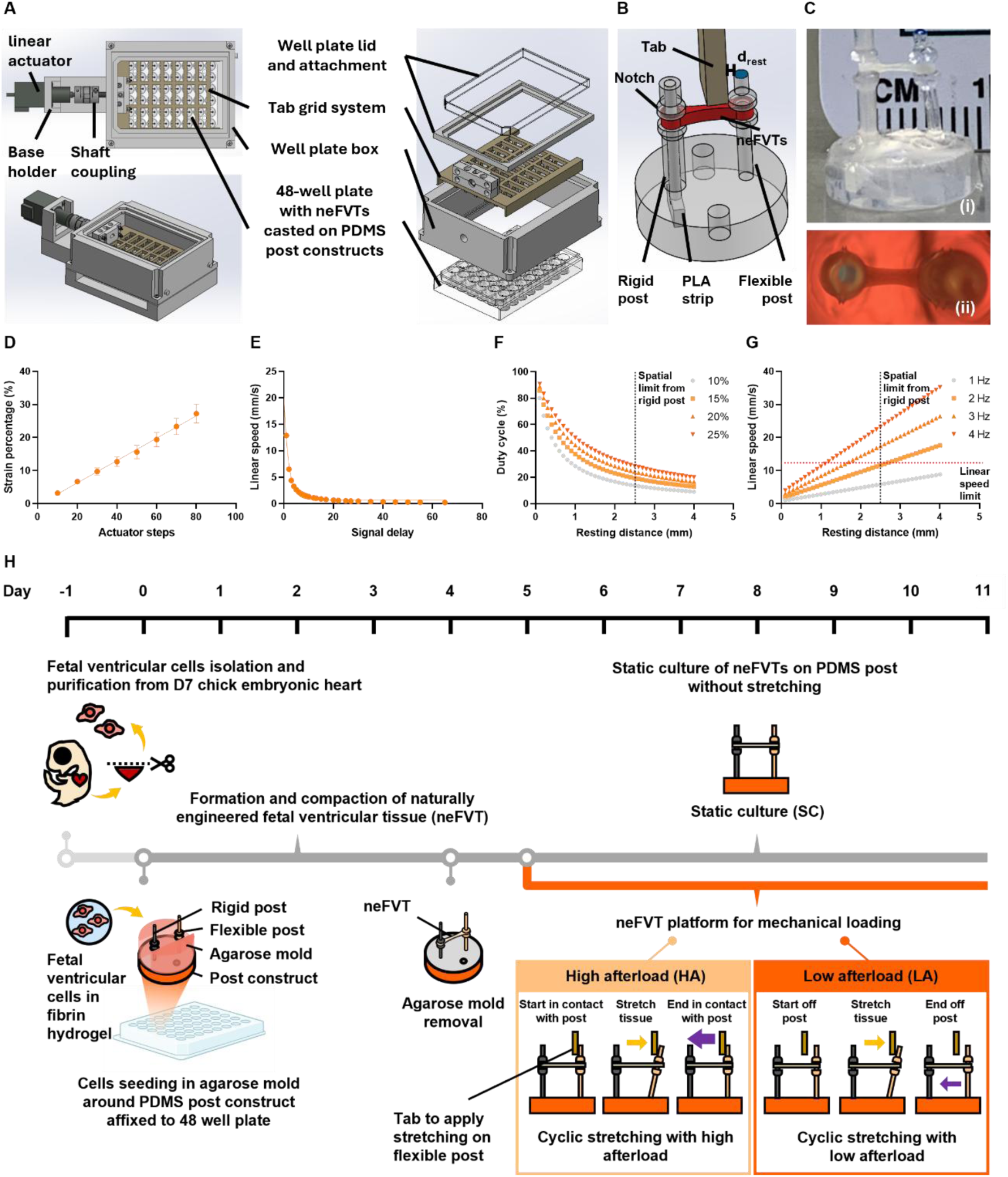
Custom, 3D printed, high throughput mechanical stimulation bioreactor allows independent modulation of afterload and asymmetrically cycled preload in NFVTs. (**A**) Configuration of the mechanical stimulation bioreactor. The bioreactor included a linear actuator connecting to a 3D printed tab grid system through a shaft coupling. The tab grid system was enclosed by a 3D printed well plate box placed over a 48 well plate containing NFVTs cast on PDMS post constructs and covered with a well plate lid secured with lid attachment. The entire assembly was mounted on a 3D printed base holder with the linear actuator. (**B**) Individual tab from the tab grid system inserted into each well of the 48 well plate to apply asymmetrically cycled preload on NFVT by pushing against the flexible post of the post construct above its notch, starting from varying resting distance (drest) away from the post. Flexible post stiffness provided afterload during tissue contraction when the tab retracted from the post and returned to its starting position designated by the resting distance. (**C**) Side and top views of NFVTs casted between the rigid and flexible posts of post construct. (**D**) Strain percentages under varying actuator steps of the linear actuator. (**E**) Linear speed under varying signal delays of the linear actuator. (**F**) Duty cycles under varying resting distances at distinct strain percentages. Black dot line indicated the spatial limit (drest < 2.5 mm) from the rigid post that restricted the distance by which the tab could retract away from the flexible post of post construct. (**G**) Linear speeds required to maintain the duty cycles at varying resting distances at 10% strain percentage. Red dot line indicated the linear speed limit (<13 mm/s) of the linear actuator. (**H**) Schematic illustration of casting NFVTs around PDMS post construct for mechanical loading. Fetal ventricular cells were first isolated from chick embryonic hearts on Day 7, purified, and rested. On Day 0, fetal ventricular cells were cast within fibrin hydrogel to form NFVT on an agarose mold around PDMS post construct in a 48 well plate. From Day 0 to Day 5, NFVT compacted into a linear tissue in between the notches on flexible and rigid posts of post construct, with agarose mold removed on Day 4. From Day 5 to Day 11, NFVTs were either continually maintained in static culture (SC), or exposed to low afterload (LA) or high afterload (HA) under asymmetrically cycled preload. Arrows represent the preload (yellow) and afterload (purple) applied on NFVT under either HA or LA. Data are means ± SEM, n > 6.

To validate the ability of individual tab in translating post deflection into tissue strain percentage at different speeds, we investigated how the input actuator variables, actuator step and signal delay, influenced the strain percentage and linear speed at the notch of flexible post. We found that the tabs reliably achieved a linear strain percentage ranging from 3% to 25% as the actuator step increased from 10 to 80, with a 5% strain error across different wells (Fig. 1D). Increasing signal delay resulted in an exponential decrease in linear speed, leading to a maximum of 13 mm/s achieved (Fig. 1E). With the experimental validation on strain percentage and linear speed, we determined the changes in duty cycle arose from varying resting distances at distinct strain percentages. Our results showed that increasing resting distance led to exponentially decreasing duty cycle, resulting in a duty cycle range from 80% down to 13% at 10% strain percentage due to the spatial limit from rigid post (d_rest_ < 2.5 mm) that restricted tab movement (Fig. 1F). Increasing strain percentage elevated duty cycle at the same resting distance as the time required to displace the flexible post increased. In this study, we performed asymmetrically cycled preload at 10% strain percentage and 16.6% duty cycle (d_rest_ = 2 mm), which facilitates the comparison with previous studies on mechanical loading of engineered heart tissue while providing ample resting distance in between stretching for tissue contractile shortening. We then determined the linear speed required to maintain the asymmetrically cycled preload at varying resting distances and frequencies under 10% strain percentage (Fig. 1G). At 1 Hz, linear speed increased from 1 to 5.8 mm/s along with the elevating resting distances until it reached the spatial limit. Increasing frequencies resulted in higher linear speed at the same resting distance that eventually exceeded the linear speed limit (< 13 mm/s) before the spatial limit. We selected 1 Hz as the stimulation frequency consistent with most previous engineered heart tissue studies employing cyclic stretching and thus a linear speed of 4.8 mm/s.

We fabricated NFVTs on PDMS post constructs in a 48-well plate format to enable integration with the mechanical stimulation bioreactor for applying distinct afterloads under asymmetrically cycled preload (Fig. 1H). Fetal ventricular cells were isolated and purified from hearts of day 7 chick embryos. Flow cytometry and immunofluorescent staining confirmed an input population of ∼80% cardiomyocytes and ∼20% cardiac fibroblasts and other cell types (Fig. S1). On day 0, cells were mixed with fibrin hydrogel and cast in an agarose mold around post constructs affixed to each well. The agarose mold constrained tissue contraction and ensured anchorage around the notches of the flexible and rigid posts during compaction. On day 4, the agarose mold was removed, yielding a compact, linear NFVT suspended between the post notches, which prevented tissue sliding during mechanical loading. Tissues that compacted with gap formation around the post notches were less responsive to mechanical stimulation and were excluded (Fig. S2). From day 5 to 11, NFVTs were either maintained in static culture (SC) or integrated with the bioreactor for application of high afterload (HA) or low afterload (LA) under asymmetrically cycled preload. In SC, NFVTs received no preload but could still contract against afterload based on the flexible post stiffness. HA applied asymmetrically cycled preload while maintaining continuous contact between the tab and the flexible post (d_rest_ = 0 mm). Under this condition, NFVTs constantly experienced effectively infinite afterload due to mechanical constraint from the tab, which prevented contractile shortening and resembled most existing engineered heart tissue models. A stretch delay was introduced following each loading cycle to maintain a 16.6% duty cycle. In contrast, LA also applied asymmetrically cycled preload but permitted contractile shortening by repositioning the tab off the flexible post after each loading cycle (d_rest_ = 2 mm). This intermittent separation reduced afterload to physiologically relevant levels (1μN/μm) and introduced a transient relaxation phase during the off-post period. As a result, LA more closely recapitulated key features of the mechanical loads within the cardiac cycle, including asymmetrically cycled preload, finite afterload, and post-contraction relaxation.

### Low afterload promotes tissue contractility with enhanced twitch kinematics

PDMS post constructs provide an intuitive means to quantify contractile force generation in NFVTs by tracking post deflection under auxotonic contraction (Fig. 2A). Under LA, NFVTs exhibited sustained contractile force generation throughout the stimulation period, leading to significantly higher normalized contractile force and contractile frequency than SC and HA as early as Day 8 that persisted through Day 11 (Fig. 2, B and C, and fig. S3). To further assess contractile function across mechanical stimulation regimens, twitch kinematics on Day 8 were analyzed (Fig. 2, D–I). NFVTs under LA exhibited significantly shorter rise time, descendent time, and total beat time compared to SC, despite generating higher forces. This resulted in increased contraction and relaxation velocities, suggesting enhanced functional maturation. To validate if the effect of LA persisted at distinct frequencies, contractile force generation of NFVTs was also assessed under LA at 2 Hz (Fig. S4). LA at 2 Hz (linear speed = 9.6 mm/s) attenuated the functional improvements observed under 1 Hz, with only minimal increases in contractile force and contractile frequency relative to SC. These findings confirm the superior performance of 1 Hz stimulation in enhancing NFVT contractile function, underscoring the importance of selecting an optimal stimulation frequency. Despite increased contractility under mechanical loading, NFVTs displayed substantial functional heterogeneity. These differential responses may stem from variations in tissue geometry and structural organization.

**Fig. 2.**
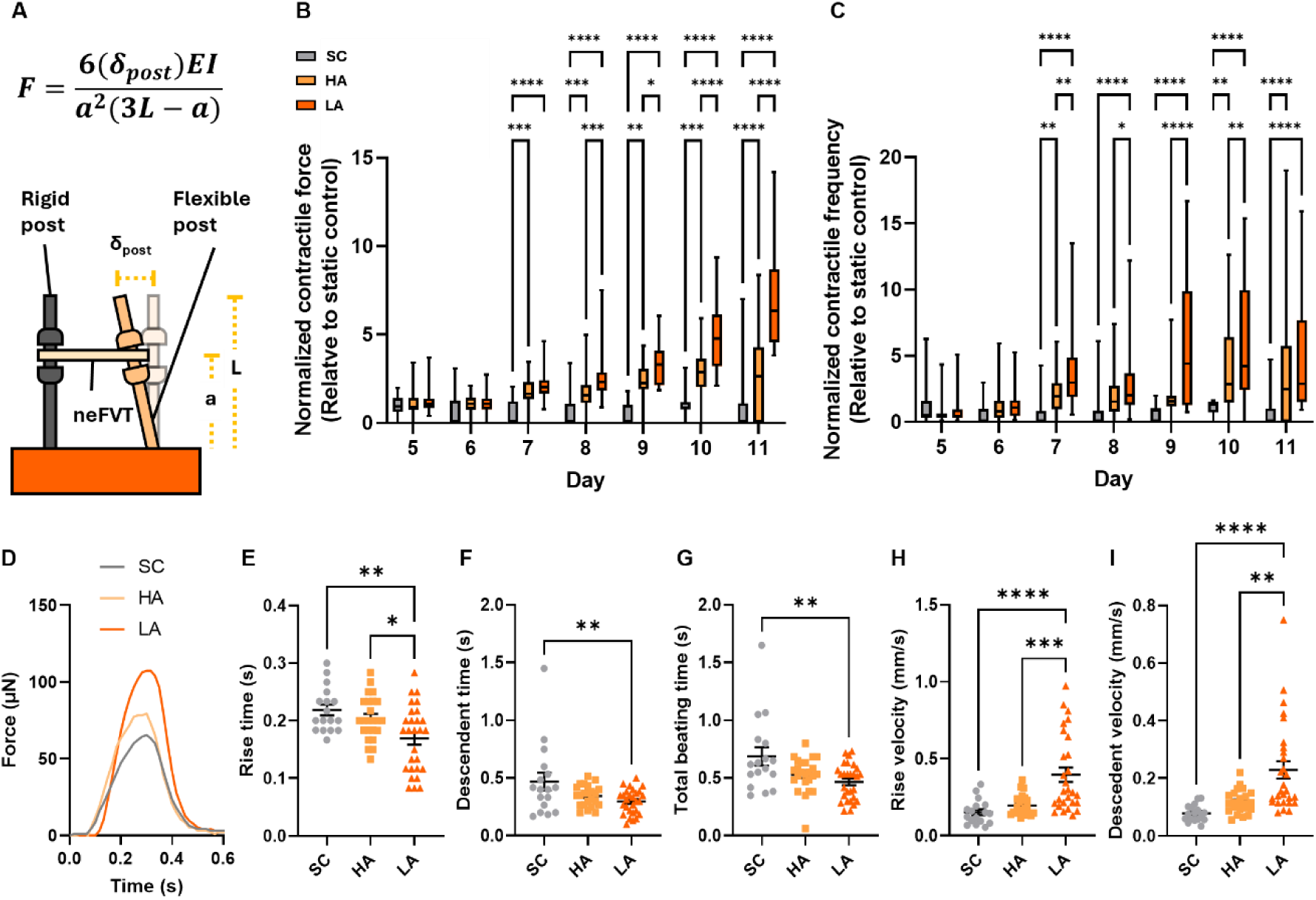
LA promotes contractile functions of NFVTs. (**A**) Schematic illustration of NFVT bending the flexible post of a post construct with force F. (**B** and **C**) Normalized contractile force (B) and frequency (C) generated NFVTs from Day 5 to Day 11 under SC, HA, and LA relative to SC. (**D**) Representative contractile force traces of NFVTs on Day 8 under SC, HA, and LA. (**E-I**) Rise time (E), descendent time (F), total beating time (G), rise velocity (H), and descendent velocity (I) of contractile force traces produced by NFVTs on Day 8 under SC, HA, and LA. Data are means ± SEM, N = 6. Two-way ANOVA with Tukey’s test; *P < 0.05, **P < 0.01, ***P < 0.001, and ****P < 0.0001.

### Low afterload facilitates sustained tissue growth associated with increasing tissue contractility

To evaluate the effects of distinct stimulation regimens on tissue growth and compaction, NFVT volume under SC, HA, and LA was derived using their top-view images on Days 5, 8, and 11 (Fig. 3A). NFVT volume was calculated as the product of tissue length between post centroids, width at the neck region, and a constant average tissue thickness of 0.3 mm derived from cryosectioning that reflects the thin nature of the NFVTs. Both HA and LA maintained significantly higher tissue volume than SC on Day 8, with preserved tissue length and width (Fig. 3, B and C, and fig. S5). However, only LA sustained tissue volume through Day 11, whereas HA exhibited a marked decline from Day 8 to Day 11. Consistently, both HA and LA reduced the percentage change in tissue volume from Day 5 to 8 relative to SC, but only LA maintained this effect from Day 8 to 11 (Fig. 3, D and E).

**Fig. 3.**
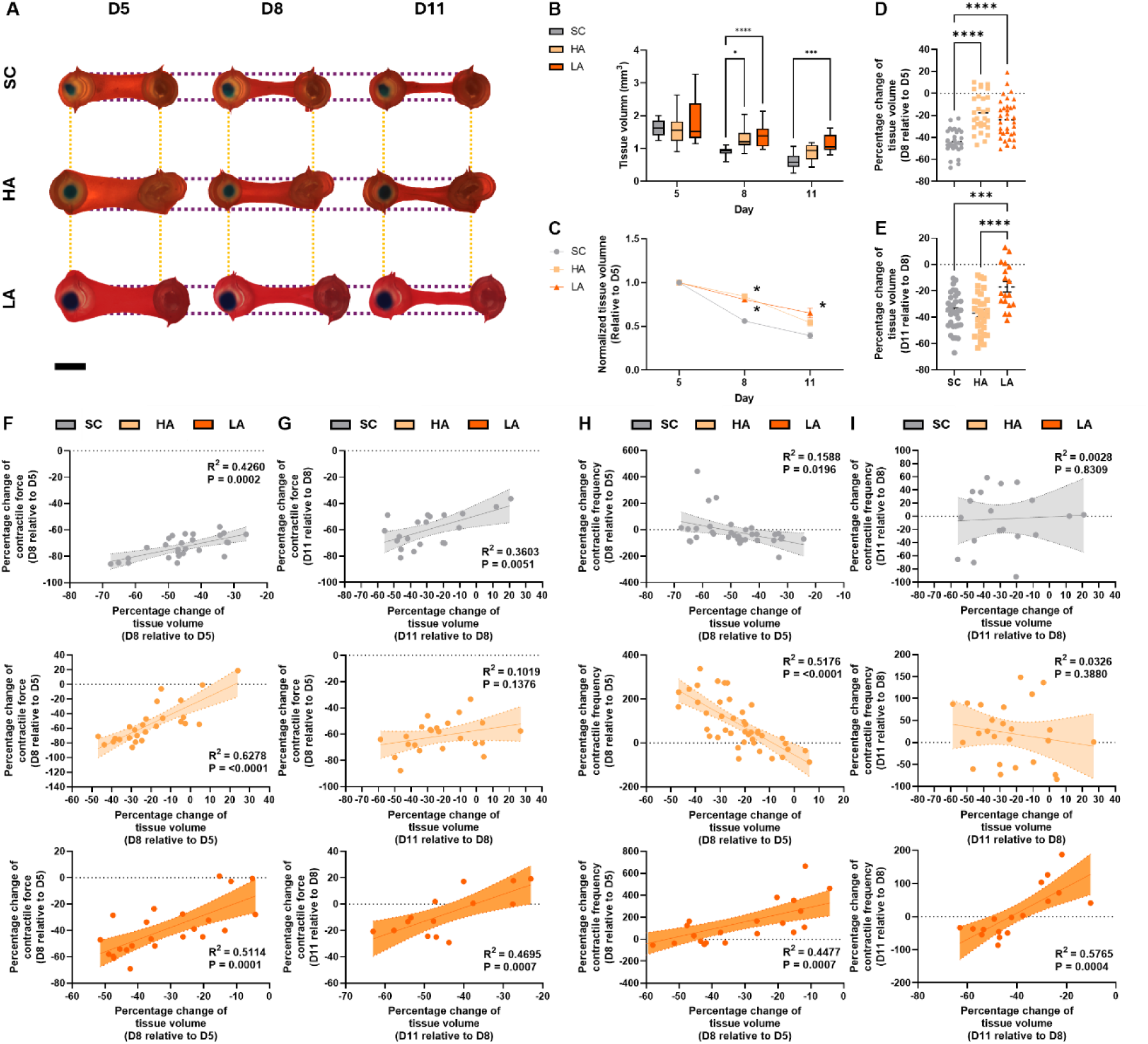
LA facilitates prolonged NFVT growth that is correlated with increasing contractile functions. (**A**) Representative top view images of NFVTs on Day 5, Day 8, and Day 11 under SC, HA, and LA. Yellow and purple dot lines indicates the initial length and width of NFVTs on Day 5, respectively. Scale bar, 2 mm. (**B** and **C**) Tissue volume (B) and normalized tissue volume relative to Day 5 (C) of NFVTs on Day 5, 8, and 11 under SC, HA, and LA. (**D** and **E**) Percentage change of tissue volume of NFVTs from Day 5 to Day 8 (D) and Day 8 to Day 11 (E) under SC, HA, and LA. (**F** and **G**) Correlation between the percentage change of tissue volume and percentage change of contractile force from Day 5 to Day 8 (F) and Day 8 to Day 11 (G) under SC, HA, and LA. (**H** and **I**) Correlation between percentage change of tissue volume and percentage change of contractile frequency from Day 5 to Day 8 (H) and Day 8 to Day 11 (I) under SC, HA, and LA. Data are means ± SEM, N = 6. Two-way ANOVA with Tukey’s test; *P < 0.05, **P < 0.01, ***P < 0.001, and ****P < 0.0001.

Correlation analysis revealed a strong positive relationship between the changes in tissue volume and contractile force from Day 5 to 8 across all groups. This relationship was weakened in SC and HA from Day 8 to 11 but remained strong in LA. Moreover, both HA and LA showed a significant correlation between the change in tissue volume and contractile frequency from Day 5 to 8, with a negative relationship for HA and a positive relationship for LA. However, only LA preserved its correlation through Day 11. Collectively, these results suggested that LA facilitates NFVTs growth throughout the stimulation period that offsets spontaneous tissue compaction and associates with increased tissue contractility

### Low afterload enhances tissue structural organization associated with cardiomyocyte arrangement, fibroblast entanglement, and tissue alignment

To investigate whether LA enhanced contractile functions of NFVTs through the improvement of structural organization, immunofluorescent staining of cardiomyocytes and cardiac fibroblasts was performed on Day 8. MF20^+^/α-SMA^-^ cardiomyocytes and MF20^-^/α-SMA^+^ cardiac fibroblasts in NFVTs revealed a dense network of cardiomyocytes ensheathed by a thin cardiac fibroblast layer, with marked differences in cellular organization and tissue alignment across mechanical loading regimens (Fig. 4A). Cardiomyocytes under LA exhibited an elongated morphology with increased cell length, resulting in a denser cardiomyocyte network compared to SC and HA (Fig. 4, B and C). Moreover, fewer cardiac fibroblasts were detected within the network, collectively suggesting the formation of a more cohesive contractile unit (Fig. 4D). Consistent with this structural feature, LA resulted in significantly greater cellular alignment than SC and HA, indicating a more anisotropic tissue structure (Fig. 4E). Notably, although the cardiomyocyte networks in SC and HA contained comparable numbers of cardiac fibroblasts on Day 8, HA resulted in a significantly higher fibroblast proportion than both SC and LA by Day 11, despite an overall decrease in fibroblast abundance across all groups (Fig. 4D, and fig. S6). This persistence of fibroblast entanglement within the cardiomyocyte network may reflect the onset of fibrotic remodeling.

**Fig. 4.**
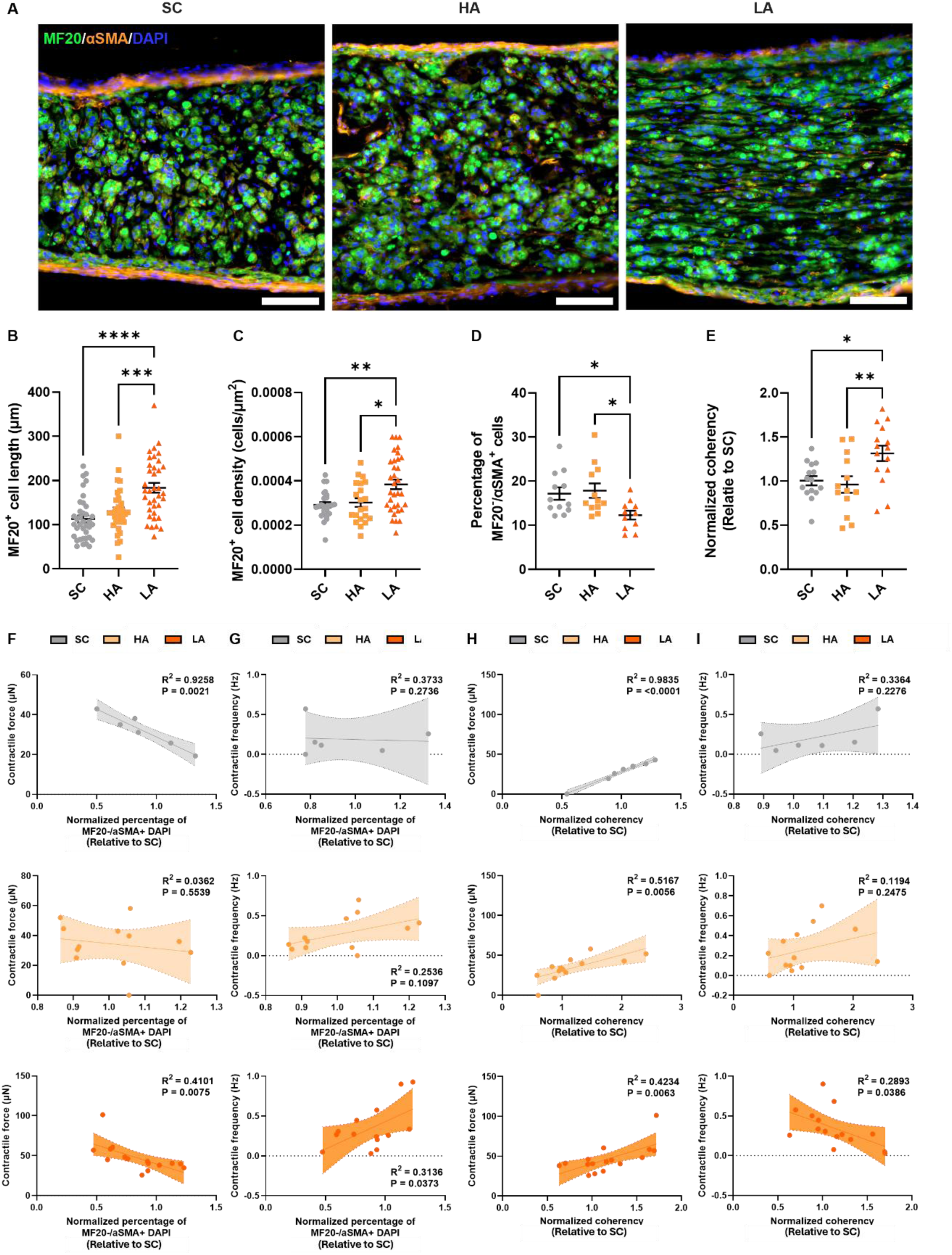
LA led to thinner and denser cardiomyocytes, reduced fibroblast entanglement and increases tissue alignment in NFVTs. (**A**) Immunofluorescent images of NFVTs stained with MF20 (green), α-SMA (orange), and DAPI (blue) on Day 8 under SC, HA, and LA. Scale bar, 200 μm. (**B**-**E**) Cardiomyocytes length (B), density (C), percentage change of MF20-/α-SMA+ cells (D), and normalized coherency (E) in NFVTs on D8 under SC, HA, and LA. (**F** and **G**) Correlation between the normalized percentage of MF20-/α-SMA+ cells on D8 and contractile force (F) or contractile frequency (G) of NFVTs from D5 to D8 under SC, HA, or LA. (**H** and **I**) Correlation between the normalized coherency on D8 and contractile force (H) or contractile frequency (I) of NFVTs from D5 to D8 under SC, HA, or LA. Data are means ± SEM, N = 6. Two-way ANOVA with Tukey’s test; *P < 0.05, **P < 0.01, ***P < 0.001, and ****P < 0.0001.

To assess whether tissue organization also predicts heterogeneous contractile function, correlations between structural metrics and contractile functions on Day 8 were analyzed. The normalized percentage of cardiac fibroblast showed a strong negative correlation with contractile force in SC and LA (Fig. 4F). In contrast, it was positively correlated with contractile frequency, reaching significance only in LA (Fig. 4G). The absence of these relationships in HA may reflect abnormal fibroblast entanglement within the cardiomyocyte network. Meanwhile, normalized tissue coherency exhibited a strong positive correlation with contractile force across all groups, but only a weak association with contractile frequency (Fig. 4, H and I). Taken together, these results indicate that LA is associated with improved structural organization of NFVTs, characterized by a denser cardiomyocyte network, reduced fibroblast entanglement, and enhanced tissue alignment, which in turn contributes to increased contractile force. More broadly, tissue structure appears to predominantly govern contractile force, with a comparatively minor influence on contractile frequency.

### Low afterload improves cardiomyocyte maturation based on sarcomeric structure, gap junction formation, gene expression, and calcium handling

To evaluate the maturation state of cardiomyocytes in NFVTs, sarcomeric structure was first examined on Day 8. Mechanical loading of NFVTs led to the formation of clear sarcomeric banding patterns, with increased Z-line width under both HA and LA (Fig. 5A). Quantitative measurements confirmed a more physiologically relevant sarcomere length (∼2.2 μm) and increased Z-line width in HA and LA relative to SC (Fig. 5, D–E). Immunofluorescent staining of the gap junction protein connexin 43 was then used to assess cell–cell electrical coupling among cardiomyocytes (Fig. 5B). Quantification revealed significantly higher connexin 43 expression in LA relative to SC and HA (Fig. 5F). Expression of the maturation marker TNNT2 was further evaluated using single-molecule fluorescent in situ hybridization (smFISH) (Fig. 5C), with upregulation observed in NFVTs under LA compared to SC and HA (Fig. 5G).

**Fig. 5.**
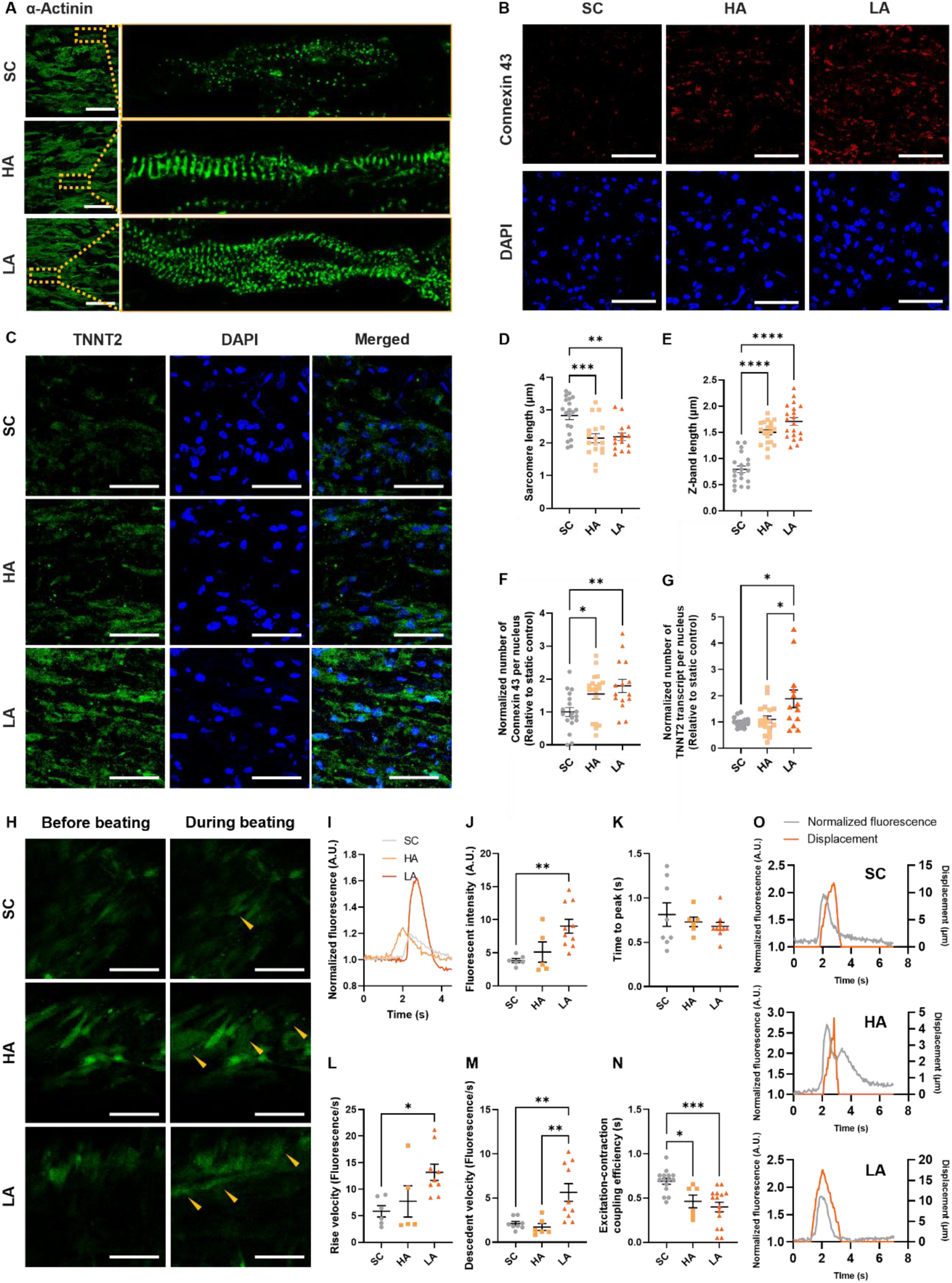
LA promotes cardiomyocyte maturation with enhanced sarcomeric organization, cell-cell interaction, transcriptomic activity, and calcium handling in NFVTs. (**A** and **B**) Immunofluorescent images of NFVTs stained with α-actinin (A) and connexin 43 (B) on Day 8 under SC, HA, and LA. Scale bars, 200 μm and 50 μm, respectively. (**C**) SmFISH images of NFVTs stained with TNNT2 on Day8 under SC, HA, and LA. Scale bar, 50 μm. (**D**-**G**) Sarcomere length (D), Z-band length (E), normalized number of connexin 43 per nucleus (F), and normalized number of TNNT2 transcript per nucleus (G) of NFVTs on Day 8 under SC, HA, and LA. (**H**) Calcium signal of NFVTs before and during beating visualized by CAL520 AM on Day 8 under SC, HA, and LA. Arrows indicate calcium spike during beating. Scale bar, 50 μm. (**I**) Representative spontaneous calcium transient of NFVTs on Day 8 under SC, HA, and LA. (**J**-**N**) Normalized fluorescent (J), time to peak (K), rise velocity (L), descendent velocity (M), and excitation-contraction coupling (N) of calcium transients in NFVT on Day 8 under SC, HA, and LA. (**O**) Representative normalized fluorescent-time curve and displacement-time curves of NFVTs under SC, HA, and LA. All data are means ± SEM, N = 3. One-way ANOVA with Tukey’s test; *P < 0.05, **P < 0.01, ***P < 0.001, and ****P < 0.0001.

To verify whether enhanced cardiomyocyte maturation in LA was accompanied by improved calcium handling, spontaneous calcium transients were recorded using CAL520 on Day 8 (Fig. 5, H and I). NFVTs under LA exhibited higher fluorescence intensity compared to SC and HA (Fig. 5J). Despite similar time to peak and rise velocity, an increased descendent velocity was observed in LA, suggesting improved calcium reuptake (Fig. 5K–M). Excitation–contraction coupling was further assessed by measuring the time from calcium transient onset to tissue displacement during contraction (Fig. 5, N and O). Mechanical loading significantly reduced excitation–contraction coupling time in both HA and LA relative to SC, with LA showing a modest additional improvement over HA.

Together, these results suggest that LA promotes a more mature cardiomyocyte phenotype in NFVTs, characterized by improved sarcomeric organization, enhanced gap junction formation, increased expression of maturation-associated genes, and more efficient calcium handling, thereby supporting the development of contractile function.

### Low afterload spares cellular turnover and suppresses fibrosis phenotypes

To investigate cellular turnover in NFVTs, we quantified apoptotic and proliferative activity by assessing cleaved caspase-3 and phospho-histone H3 (pHH3) expression, respectively (Fig. 6, A and B). NFVTs under LA exhibited minimal cleaved caspase-3 expression, comparable to SC, whereas HA significantly increased cleaved caspase-3 expression, indicative of enhanced apoptotic activity (Fig. 6C). Analysis of pHH3 expression revealed increased proliferative activity in both HA and LA relative to SC, although statistical significance was reached only in HA (Fig. 6D). Together, these findings suggest elevated cellular turnover under HA, characterized by concurrent increases in apoptosis and proliferation, whereas LA preserved cellular turnover at levels comparable to SC. To determine whether the altered cellular turnover observed under HA was associated with pathological remodeling, we next examined the expression of fibrosis-associated genes using single-molecule fluorescence in situ hybridization (smFISH) on Day 8 NFVTs (Fig. 6, E and F). HA significantly upregulated the expression of POSTN, TGFB2, and COL5A2 compared with SC, whereas expression levels under LA remained similar to those observed in SC (Fig. 6, G–J). Expression of FN1 showed no significant differences among groups (Fig. 6H). These data indicate that HA promotes a profibrotic transcriptional program in NFVTs, whereas LA largely preserves a non-fibrotic phenotype. Collectively, these results demonstrate that HA is associated with increased cellular turnover and induction of fibrosis phenotypes in NFVTs. In contrast, LA maintains cellular homeostasis and limits fibrotic remodeling, supporting tissue maturation while suppressing pathological remodeling.

**Fig. 6.**
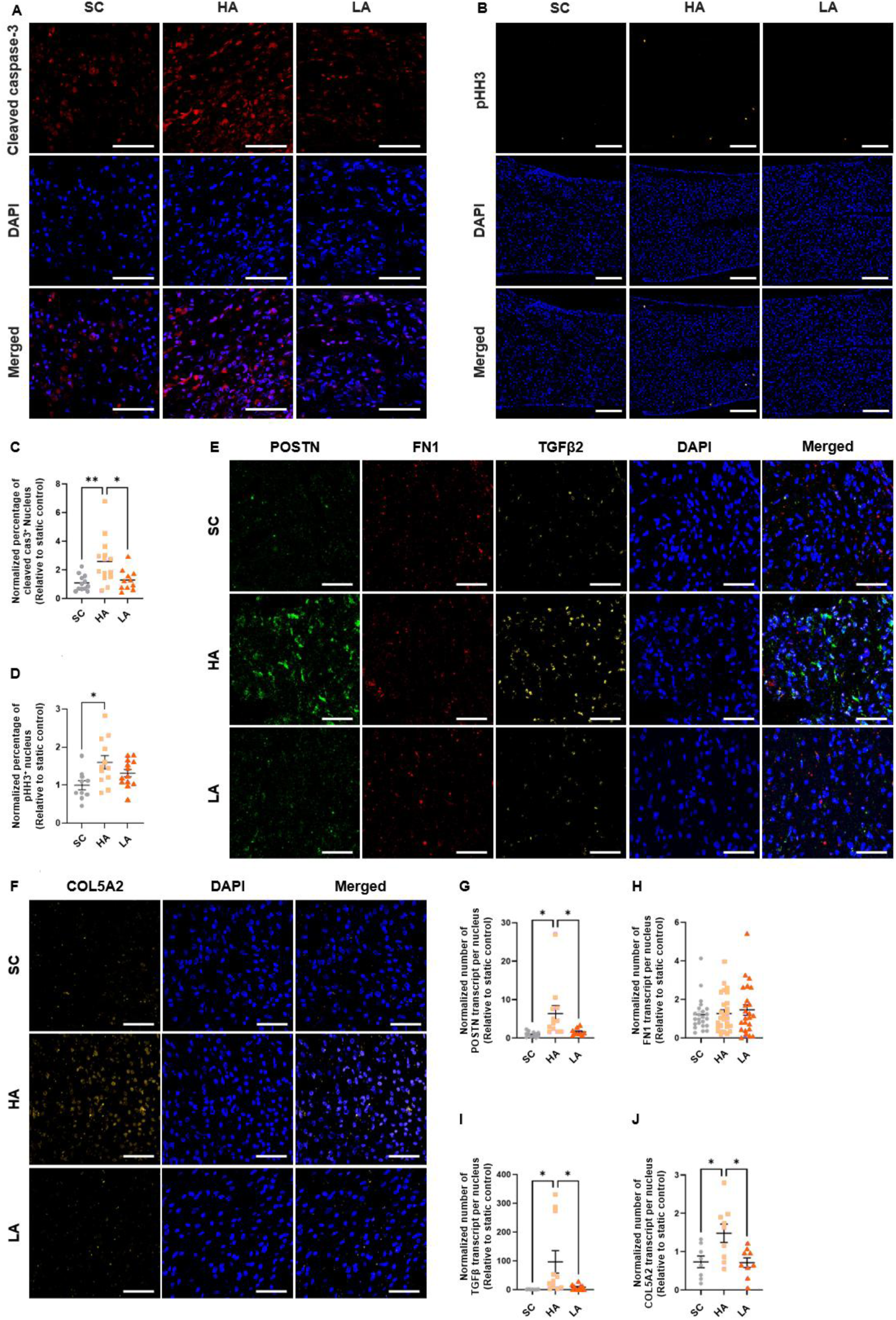
LA spares cell turnover and suppresses fibrosis phenotypes in NFVTs. (**A** and **B**) Immunofluorescent images of NFVTs stained with cleaved caspase-3 (A) and pHH3 (B) in NFVTs on Day 8 under SC, HA, and LA. Scale bars, 50 μm and 200 μm, respectively. (**C** and **D**) Normalized percentage of cleaved caspase-3+ (C) and pHH3+ (D) nucleus relative to SC in NFVTs on Day 8 under SC, HA, and LA. (**E** and **F**) smFISH images of NFVTs stained with POSTN, FN1, TGFβ2 (E), and COL5A2 (F) on Day 8 under SC, HA, and LA. Scale bar, 50 μm. (**G**-**J**) Normalized number of POSTN (G), FN1 (H), TGFβ2 (I), and COL5A2 (J) transcripts per nucleus relative to SC in NFVTs on Day 8 under of SC, HA, and LA. All data are means ± SEM, N = 3. One-way ANOVA with Tukey’s test; *P < 0.05, **P < 0.01, ***P < 0.001, and ****P < 0.0001.

### Low afterload maintains minimal PIEZO1 expression and inhibits YAP and NOTCH activation in cardiomyocytes

We sought to elucidate the molecular mechanisms by which mechanical loading modulates NFVT maturation or pathogenesis through cardiomyocyte mechanotransduction. We focused on the mechanotransducer PIEZO1, YAP1, and NOTCH and performed smFISH and immunofluorescent staining together with the cardiomyocyte marker MF20 (Fig. 7, A–C). HA markedly increased PIEZO1 transcript levels, with the majority localized to MF20^+^ cardiomyocytes (Fig. 7A). In contrast, SC and LA exhibited minimal PIEZO1 expression. Quantification confirmed significantly elevated PIEZO1 transcripts in HA relative to both SC and LA, suggesting cardiomyocyte PIEZO1 overexpression (Fig. 7D). Consistent with increased PIEZO1 expression, HA promoted YAP1 nuclear localization in MF20^+^ cardiomyocytes, whereas LA showed reduced nuclear YAP1 (Fig. 7B). Quantification of nuclear YAP1 levels confirmed higher expression in HA relative to SC and reduced expression in LA (Fig. 7E). These findings suggest activation of PIEZO1–YAP1 signaling in cardiomyocytes under HA. In contrast to YAP1, activated NOTCH expression in HA was comparable to SC but significantly reduced in LA (Fig. 7B). Quantification of nuclear activated NOTCH levels in MF20^+^ cardiomyocytes showed a modest decrease in HA and a pronounced reduction in LA relative to SC (Fig. 7F).

**Fig. 7.**
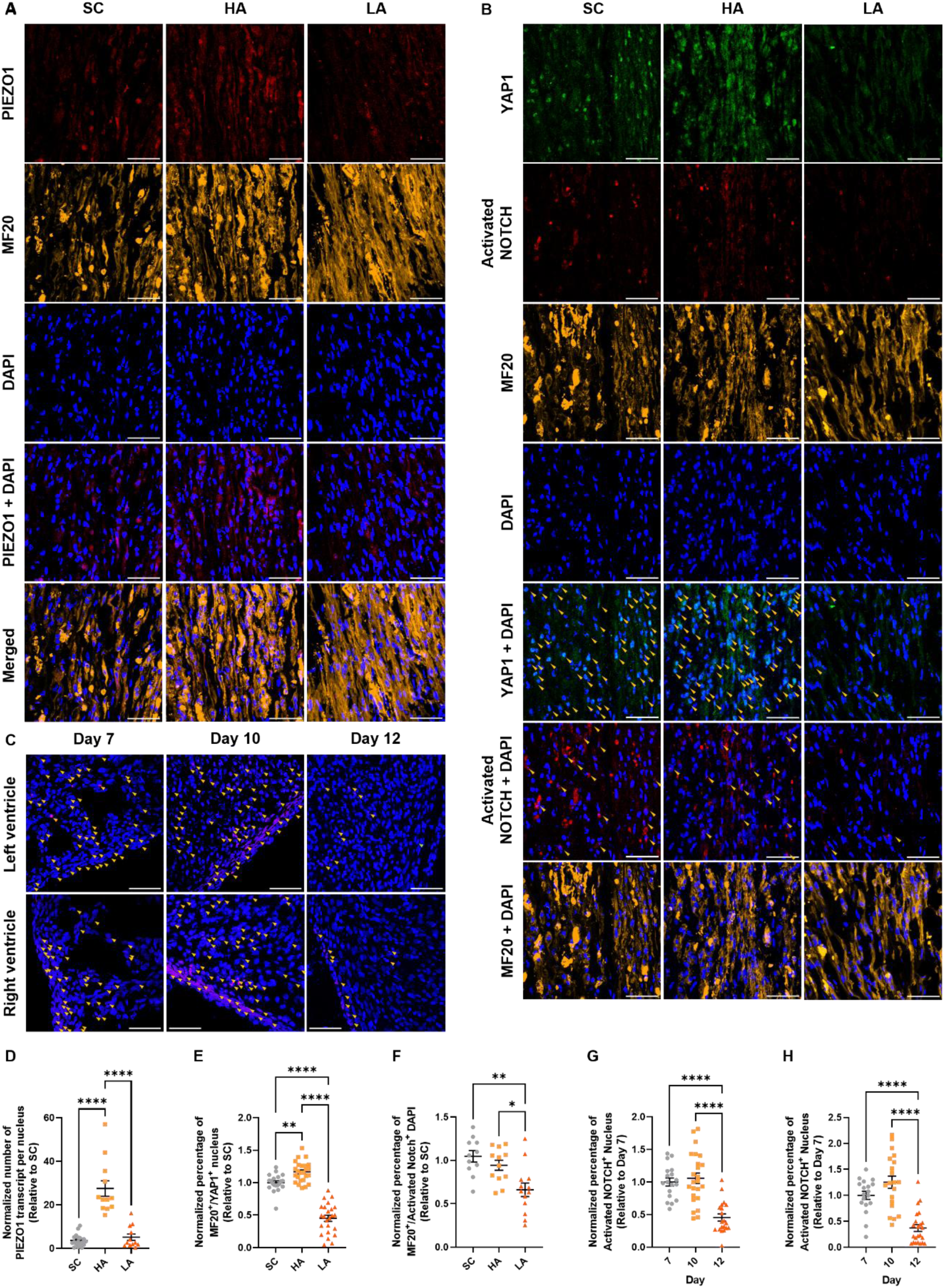
LA inhibits PIEZO1 downstream targets YAP and NOTCH in cardiomyocytes within NFVTs. (**A**) Images of smFISH staining with PIEZO1 and immunofluorescent staining with MF20 in NFVTs on Day 8 under SC, HA, and LA. Scale bar, 50 μm. (**B**) Immunofluorescent images of NFVTs stained with YAP1, activated NOTCH, and MF20 on Day 8 under SC, HA, and LA. Scale bar, 50 μm. (**C**) Immunofluorescent images of left and right ventricles in chick embryonic heart section stained with activated NOTCH on Day 7, Day 10, and Day 12. Scale bar, 50 μm. (**D**-**F**) Normalized number of PIEZO1 transcripts per nucleus (D), normalized percentage of MF20+/YAP1+ nucleus (E), and normalized percentage of MF20+/activated NOTCH+ nucleus (F) relative to SC in NFVTs on Day 8 under SC, HA, and LA. (**G** and **H**) Normalized percentage of activated NOTCH+ nucleus relative to Day 7 in left (G) and right (H) ventricles of chick embryonic heart section on Day 7, Day 10, and Day 12. All data are means ± SEM, N = 3. One-way ANOVA with Tukey’s test; *P < 0.05, **P < 0.01, ***P < 0.001, and ****P < 0.0001.

To determine whether suppression of YAP1 and NOTCH signaling under LA reflects native developmental programs, we assessed activated NOTCH expression in chick ventricular myocardium at embryonic Days 7, 10, and 12 (Fig. 7C). Activated NOTCH expression was sustained from Day 7 to Day 10 but decreased by Day 12 in both left and right ventricles. Quantification confirmed a significant reduction on Day 12 relative to Days 7 and 10 (Fig. 7, G and H). Taken together, these results demonstrate that LA maintains minimal PIEZO1 signaling and downregulates YAP1 and NOTCH, recapitulating aspects of normal myocardial development. In contrast, HA potentially activates PIEZO1–YAP1 signaling, which may contribute to the pathological responses observed under high afterload.

## Discussion

This study reports the development of NFVT platform for elucidating the mechanisms by which the natural cardiac loading condition regulates growth, maturation, and pathological remodeling of the developing myocardium at the cellular and mesoscale level. Our results support that LA is essential to augmented contractile function of NFVT with sustained tissue growth, enhanced cardiomyocyte maturation, and suppressed fibrosis phenotypes. These phenotypic changes were mediated by the regulation of cardiomyocyte mechanotransduction, including minimal PIEZO1 expression and suppressed YAP1 and NOTCH activation, and the enhancement of tissue architecture, characterized by increased tissue alignment and reduced fibroblast entanglement. In contrast, HA led to functional dysfunction accompanied by fibrosis remodeling potentially bought by the activation of cardiomyocyte PIEZO1/YAP1 signaling and maintained fibroblast entanglement.

The NFVT platform is built based on knowledge gained from previous engineered heart tissue models for mechanical loading using primary, hESC-, and hiPSC-derived cardiomyocytes. However, the key issue of most of these models lies in the restriction of contractile shortening during mechanical loading, which coupled preload with effectively infinite afterload that impedes normal tissue maturation and induces a disease state within engineered heart tissues (17–19,21,24,31,32). Recent studies have focused on the development of novel systems that are capable of independently and temporally modulating preload and afterload while facilitating tissue shortening (15,25). Nonetheless, these systems are limited to passive, stepwise preload and lack capacity to fully mimic other fundamental features that characterize dynamic preload in the native heart, including frequency, pattern, and duty cycle. These can greatly reduce their physiological relevance and hinder the recapitulation of morphogenic and mechanistic events underlying myocardial development driven by mechanical loadings.

We demonstrated that LA achieved sustained tissue contractility in NFVTs with enhanced twitch kinetics, resulting in higher contractile force and frequency compared to SC and HA under auxotonic conditions. This contrasts with most previous findings showing that increasing afterload correlates with progressively higher contractile force (23,25,26,28). This apparent discrepancy likely reflects differences in the afterload range being compared, in which HA represents an effectively infinite (rigid) afterload that fully restricts contractile shortening, whereas most prior studies examine increases within a finite afterload range that still permits shortening. It has also been reported that prolonged culture of engineered heart tissue at high afterload led to loss of contractile force compared to those cultured at moderate afterload, supporting our observation that NFVTs under effectively infinite afterload exhibited reduced contractile force generation (24). Interestingly, consistent with prior observation of engineered heart tissue under low afterload, NFVTs exposed to LA exhibited higher rise and descendent velocity of the contractile force traces compared to SC and HA (23). Despite the overall improvement in contractile performance, we observed substantial variability in tissue contractility under the same mechanical loading regimen. This variability was attributed to the differences in individual tissue geometry and structural organization, with our data showing that tissue contractility strongly depends on tissue geometry and structural organization.

We further demonstrated that LA is required to sustain NFVT growth with improved contractile function. Compared to SC and HA, NFVTs under LA maintained tissue volume despite ongoing compaction, resulting in persistent associations with enhanced contractile force and frequency throughout the stimulation period. Preservation of tissue volume under LA is consistent with continued tissue growth, a hallmark of the developing myocardium. During cardiac development, myocardial growth is initially driven by cardiomyocyte hyperplasia during the prenatal period and subsequently supported by physiological hypertrophy after birth (33,34). Consistent with this developmental paradigm, NFVTs exposed to LA exhibited increased cardiomyocyte density, elevated proliferative activity, and minimal apoptosis, collectively suggesting active tissue growth supported by cardiomyocyte and non-cardiomyocyte expansion with favorable cell survival. In contrast, the marked reduction in tissue volume observed in NFVTs under HA by Day 11 occurred despite increased proliferative activity and may reflect a net loss of tissue resulting from elevated apoptosis.

Cardiomyocyte maturation is commonly characterized by progressive improvements in its morphology and structural organization, including cell shape, alignment, and sarcomeric organization (35,36). We showed that LA led to cardiomyocyte elongation with increased density and alignment, resembling cellular events during myocardial compaction and later during early postnatal cardiac development (6,37). Moreover, NFVTs exposed to LA exhibited a physiological working range of sarcomere from 1.8 to 2.2 μm consistent with prior engineered heart tissue studies (17,18,38,39). In addition to morphological and structural maturation, we assessed the maturity of cardiomyocyte through the expression of protein and gene maturation markers (40). Connexin 43, a key gap junction protein involved in intercellular electrical coupling, is one of the most commonly used markers for this purpose. Several studies using hiPSC-derived cardiomyocytes have reported decreased or unchanged Cx43 expression following mechanical stimulation, potentially due to the application of relatively low strain amplitudes and the intrinsic 2D nature of the platforms (31,41). In contrast, we observed an approximately 2.0-fold increase in Cx43 expression across all loading regimens, consistent with studies employing a 10% input strain (19,42,43). Furthermore, we evaluated the transcriptional activity of TNNT2 gene, which encodes cardiac troponin T (cTnT), a core sarcomeric protein essential for force generation. Cardiomyocyte maturation is accompanied by an upregulated TNNT2 expression to support the assembly and organization of sarcomere during development (31,38,44,45). Accordingly, we showed that LA-conditioned NFVTs exhibited upregulated TNNT2 expression, in agreement with previous studies correlating elevated cTnT expression to enhanced cardiomyocyte maturation. Lastly, cardiomyocyte maturation was further assessed by measuring calcium handling properties of NFVTs. LA significantly enhanced calcium handling, as evidenced by increased calcium transient rise and decay velocities and shortened excitation–contraction coupling times. Together, these data confirmed that LA, which faithfully resembled the natural cardiac loading condition, possessed superior capacity to promote cardiomyocyte maturation in NFVTs, thereby leading to improved tissue functionality.

The ability to independently modulate afterload and asymmetrically cycled preload in NFVTs not only opens up opportunities to better understand the role of natural cardiac loads in normal myocardial development, but also shed light on congenital heart disease that arises from defective myocardial growth and maturation caused by abnormal loading conditions. In this study, we showed that HA resulted in contractile dysfunction in NFVTs that was attributed to insufficient tissue growth, compromised cardiomyocyte maturation, and emergence of fibrotic remodeling. Specifically, NFVTs under HA exhibited shortened growth, reduced TNNT2 transcript expression, elevated transcriptional activity of fibrosis-associated genes, and persistent cardiac fibroblast entanglement compared to LA. Altered hemodynamic loading has long been recognized as a driver of cardiac defects that resemble those observed in congenital heart disease (46,47). Although the etiology of congenital heart disease is multifactorial and remains an active area of investigation, several pathological features commonly characterize disease progression, including impaired myocardial growth, disrupted cardiomyocyte maturation, and cardiac fibrosis. For example, the underdeveloped left ventricle in hypoplastic left heart syndrome reflects abnormal myocardial development characterized by compromised growth and maturation, ultimately resulting in a thinner ventricular wall (48–50). In addition, reduced expression of contractile and maturation-associated genes has been reported in cardiomyocytes from patients with pulmonary atresia with intact ventricular septum and ventricular septal defect (49–51). Cardiac fibrosis, defined by the excessive accumulation of fibrillar collagen within the myocardial interstitium, is also frequently observed in congenital heart diseases associated with abnormal mechanical loading, including tetralogy of Fallot, the systemic right ventricle, and following arterial switch operations (51,52). Taken together, these findings suggest that NFVTs subjected to HA recapitulate key aspects of stress-induced myocardial remodeling associated with congenital heart disease that are difficult to study in vivo. Previous studies have reported pathological hypertrophy, fibrosis, and a shift toward fetal-like metabolism in engineered heart tissue models exposed to elevated afterload using neonatal rat ventricular cells or human induced pluripotent stem cell-derived cardiomyocytes (23,24). However, these models lack active preload and therefore do not fully reproduce the physiological loading pattern and duty cycle experienced by the developing myocardium. Consequently, their ability to capture the full spectrum of pathological responses associated with abnormal developmental loading

Mechanotransduction is a fundamental regulator of both adaptive and maladaptive myocardial remodeling in response to mechanical stress. Among known mechanotransducers, PIEZO1 is a mechanosensitive cation channel that mediates calcium influx and downstream signaling upon mechanical stimulation (53,54). In endothelial and epithelial cells, PIEZO1 is essential for stem cell differentiation, vascular development, and valvulogenesis during cardiac morphogenesis (55,56). In contrast, aberrant PIEZO1 activation in cardiomyocytes disrupts calcium and lipid homeostasis, promoting hypertrophy and adverse remodeling in mechanically overloaded adult mouse and human hearts (57–59). These pathological effects have been linked to downstream modulation of YAP1- and NOTCH-associated signaling pathways (60,61). Consistent with these findings, HA induced marked upregulation of PIEZO1 expression and increased nuclear localization of YAP1 in cardiomyocytes relative to SC controls, indicating activation of the PIEZO1–YAP1 mechanotransductive axis. The concomitant emergence of fibrotic phenotypes under HA further supports a maladaptive role for PIEZO1 signaling in the fetal myocardium, analogous to its established function in adult cardiac remodeling. These observations may explain the pathological remodeling previously reported in engineered heart tissue platforms exposed to effectively infinite afterload. Meanwhile, NFVTs subjected to LA maintained minimal PIEZO1 expression comparable to SC and exhibited reduced YAP1 activity alongside enhanced NOTCH activation. This signaling profile more closely recapitulates the physiological developmental maturation program of the native myocardium. Previous studies have demonstrated that YAP1 and NOTCH signaling sustain proliferative and immature cardiomyocyte states (62–64), whereas suppression of these pathways is required for initiation of cardiomyocyte maturation programs(65,66). Collectively, these findings demonstrate that mechanical loading alone can refine the developmental signaling networks governing myocardial maturation, achieving effects not attainable in previous engineering heart tissue models employing active cyclic stretch.

As an evolving experimental platform, the current mechanical stimulation bioreactor system presents opportunities for continued improvement. To commence with, there was a lack of precise control over the ratio of input cell populations. Although the overall ratio of cardiomyocyte and cardiac fibroblast (4:1) following cardiomyocyte purification was quantified, this ratio could vary among different experimental replicates and led to heterogeneous NFVT responses following mechanical loading. A lower cardiomyocyte to cardiac fibroblast ratio (7:3) may also improve NFVT functionalities (67). Furthermore, the bending stiffness of flexible post on the PDMS post construct was determined by the base-to-curing agent mixing ratio during PDMS post construct fabrication. Therefore, despite the opportunity to customize the magnitude of afterload, the current system did not currently support the adjustment of afterload during the stimulation period upon casting of NFVT on the post construct. Future incorporation of a piezoelectric system that allows the anchorage of NFVT while supporting active control of post effective stiffness could further expand the ability of the current bioreactor system to capture the physiological loading conditions. Last but not least, fixed loading freqency and phases were employed in the loading regimen presented here to enforce spontaneous contraction of NFVT following each active stretching in LA. However, it was still possible that NFVT contraction occurred during active stretching when afterload remain effectively infinite.

In conclusion, we have developed a platform for the biomimetic loading of NFVTs using distinct afterloads under asymmetrically cycled preload, which closely resemble the natural cardiac loading environment. By tuning afterload in NFVTs, we were able to either improve tissue functions with simultaneous growth and maturation in LA or induce functional impairment with compromised maturation and pathological remodeling in HA. We also highlighted the fundamental role of cellular level cardiomyocyte mechanotransduction and mesoscale tissue architecture in mediating myocardial growth, maturation, and disease.

## Materials and Methods

### Experimental design

The objective of the current study was to develop a naturally engineered fetal ventricular tissue (NFVT) platform for (i) the application of distinct afterloads under asymmetrically cycled preload on NFVTs using a custom, 3D printed mechanical stimulation bioreactor, (ii) assessment of NFVT contractile function and phenotypes, and (iii) investigation of cellular signaling and structural remodeling within NFVT. We designed the mechanical stimulation bioreactor primarily based on the combination of 3D printed tab grid system and PDMS post construct in a 48-well plate setting. We examined NFVTs subjected to static culture and low and high afterload using contractility and geometry analysis, immunochemistry, single molecule fluorescence in situ hybridization, calcium imaging, and electron microscopy. In this study, fetal ventricular cells were isolated from the hearts of day 7 chick embryos. All experimental procedures were approved by Cornell University Institutional Animal Care and Use Committee (IACUC) and were exempt from full IACUC review as the use of chick embryos at early developmental stages was not considered live vertebrate animal under Public Health Service (PHS) policy. A minimum of three independent biological replicates were obtained from multiple experiments and used to confirm the data.

### Fabrication of mechanical stimulation bioreactor with control system

The mechanical stimulation bioreactor presented here was designed in SolidWorks. All plastic components were 3D printed with polyethylene terephthalate glycol (PETG) using an Object Pro 3D printer. Tab grid system was 3D printed with polypropylene (PP) via multijet fusion 3D printing (Sculpteo). Nema 11 captive linear actuator motor (Nanotec, LGA281S10-A-UGFC-019) was used to provide a linear translation of 0.0254 mm/step to drive the tab grid system via a clamping flexible shaft coupling (McMaster-Carr, 2401K15) for tunable uniaxial cyclic stretch. The components that were designed to come into close contact with NFVTs, including the well plate box, well plate attachment, and tab grid system, were assembled and sterilized via incubation with sterile filtered acidic 70% (v/v) ethanol (pH 2.0) in distill water for at least 2 hours at room temperature and washed with autoclaved 18 MΩ·cm water.

The captive linear actuator motor was controlled by a custom Arduino Uno circuit. The circuit involved an Arduino Uno that communicated with an A4988 microcontroller to adjust the linear speed, directionality, and linear displacement of the actuator motor. A custom Arduino Uno program was developed to allow user inputs for achieving desired mechanical stimulation regimen to load the NFVTs. The entire circuit setup was connected to the linear actuator with long wires such that the circuit could be maintained at room temperature during mechanical loading of NFVTs under incubator environment.

### Fabrication of PDMS post construct in 48-well plate

PDMS post constructs were created by casting SYLGARD 184 PDMS (Dow Corning) at a 10:1 base–to–curing agent ratio in a base mold inserted with two-part sleeve components for post geometry. The base mold and two-part sleeve components were designed in SolidWorks and 3D printed with nylon PA12 using a selective laser sintering 3D printer (Sculpteo) and veroclear using an Object Pro 3D printer, respectively. The mold with PDMS was degassed under vacuum for at least 30 min and T-shape rods 3D printed with PLA using an Ender 3 3D printer was inserted into one of the sleeve components assembly to create a PDMS encapsulated rigid post. PDMS post constructs were cured at 65 ℃ for at least 1 hour and removed from the mold. All constructs were sterilized by autoclaving following the installation of metal pins (McMaster-Carr, 91585A909).

Sterilized PDMS post constructs were attached to the bottom of each wells in a 48-well plate (Greiner Bio-One, 677102) using uncured SYLGARD 184 in a 10:1 base–to–curing agent ratio as an adhesive. The PDMS post constructs were aligned across the well plate by inserting into alignment tools designed in SolidWorks and 3D printed with veroclear using an Objet Pro 3D printer via metal pins and assembling with the 48-well plate. The well plates were then incubated at 65 ℃ for at least 1 hour together with alignment tools to cure the PDMS adhesive, sterilized via incubation with acidic 70% ethanol for at least 1 hour at room temperature following removal of alignment tools, and washed with autoclaved 18 MΩ·cm water.

### Chick fetal ventricular cells isolation and purification

Chick hearts were obtained from fertile Bovans brown eggs purchased from a local farmer and incubated in a standard temperature- and humidity-regulated incubator for 7 days (Hamburger-Hamilton stage 31). The use of chick embryos at this developmental stages does not require ethical board approval. Ventricular heart tissues were dissected from chick heart and stored temporally in HBSS with calcium and magnesium (Gibco, 14025092). Isolated ventricular heart tissues were dissected into small fragments and digested by two 10 min treatments with 0.05% (w/v) trypsin (Sigma, T7409) in HBSS without calcium and magnesium (Gibco, 14170120) at 37 ℃. Embryonic ventricular cells were dissociated from the digested tissues by four 5 min treatments with 0.05% (w/v) trypsin in HBSS without calcium and magnesium at 37 ℃. Dissociated cells in trypsin obtained following each 5 min treatment were filtered through a 70 μm cell strainer, followed by an equal volume of Leibovitz’s L15 media (Gibco, 11415064), supplemented with 10% (v/v) fetal bovine serum (FBS, GeminiBio, S11150) and 1% (v/v) penicillin streptomycin. All cells were plated onto standard cell culture dishes at a density of 30,000 cells/cm^2^ in M199 media without L-glutamine (Sigma, M2154), supplemented with 3% (v/v) FBS and 1% (v/v) penicillin/streptomycin overnight.

Chick fetal ventricular cells were purified to achieve the optimal cardiomyocyte-to-cardiac fibroblast ratio prior to ventricular engineered heart tissues (NFVTs) fabrication. Chick embryonic ventricular cells plated overnight were detached and plated onto standard cell culture dishes at a density of 30,000 cells/cm^2^ in M199 media with L-glutamine (Gibco, 11150059), supplemented with 10% (v/v) FBS and 1% (v/v) penicillin/streptomycin at 37 ℃ for 45 min. Cell suspension in the cell culture dishes was collected following one wash using the culture media.

### Flow cytometry

Purified chick fetal ventricular cells suspension was fixed in 4% (v/v) formaldehyde (Electron Microscope Sciences, 15710) in FACS buffer constituted of 1X PBS, 2% FBS, and 0.02 % sodium azide (Sigma, S2002) for 20 min and washed with FACS buffer at room temperature twice. FACS buffer was pre-chilled at -20 °C prior to use. Cell suspension was then permeabilized with 0.1% (v/v) Triton-X 100 (Sigma, X100) in FACS buffer for 20 min and washed with FACS buffer at room temperature twice. 0.5 mg/ml MF20 antibody (Invitrogen, 14-6503-82) and DAPI (Invitrogen, R37606) in FACS buffer were then pipetted to the cell suspension for a 1 hour incubation at room temperature. Following washes in FACS buffer thrice, the cell suspension was incubated with Alexa Fluor Plus 488 (Invitrogen) at 1:500 dilution in FACS buffer for 30 min at room temperature. Cell suspension was washed and resuspended in FACS buffer for analysis on flow cytometer.

All samples were acquired using an Attune NXT cytometer (Thermo Fisher Scientific). Flow cytometry acquisitions were analyzed using Attune NXT cytometric software. A minimum of 20,000 events were collected in each experiment.

### Fabrication of naturally engineered fetal ventricular tissue

Before NFVTs fabrication, agarose mold was cast around the PDMS post construct. A two-part agarose mold template designed in SolidWorks and 3D printed with PLA using an Ender 3 3D printer was inserted into the 48-well plate with PDMS post constructs. 1% (w/v) agarose type II (VWR, 0815-100G) in 1X M199 without L-glutamine (Sigma, M0650) was cast around the agarose template by allowing it to gel at -20 ℃ for 20 min. Agarose residue was removed using cell culture vacuum pump after solidification of the agarose mold and removing the agarose template.

NFVTs were created by embedding purified chick embryonic ventricular cells into fibrin hydrogel with a final concentration of 10 mg/ml fibrinogen (Sigma, F8630) in 1X PBS (Gibco, 10010023), 50 U/ml thrombin (Sigma, T4648) in 1X PBS with 0.1% (w/v) bovine serum albumin (BSA, Sigma, A7030), 20% (v/v) 5X M199 without L-glutamine supplemented with sodium bicarbonate, 3% (v/v) FBS, 55.4% v/v autoclaved 18 MΩ·cm water, and 4 X 10^6^ total cells/ml consisting of 80% cardiomyocytes and 20% cardiac fibroblasts and other cells. Ingredients were mixed with a pipette, and then 100 μm of the cell mixture was pipetted into agarose molds in each well around the PDMS post construct. The well plates containing agarose molds and cell suspension around PDMS post constructs were then placed at 37 °C to allow for gelation. After a 20 min incubation period, NFVTs were incubated and maintained in M199 media without L-glutamine supplemented with 3% FBS and 1% penicillin/streptomycin.

### Tissue contractility measurement

48-well plate containing NFVTs cast on PDMS post construct was disassembled from the bioreactor system and equilibrated under static culture for 30 min prior to imaging. NFVTs were imaged with an Iphone XR mounted onto a Dual Lit Halogen Trinocular Stereo Zoom Microscope (Amscope) via an Iphone microscope adapter (LabCam) to record the 2D translation of flexible post during tissue contraction. The microscope provided a heat stage to maintain a proper culture temperature during imaging. Custom ImageJ and MATLAB programs were created to automatically calculate the change in length, contractile force, and contractile frequency of tissues from contraction videos.

### Immunofluorescence

Tissues were fixed in 4% formaldehyde overnight at 4 °C, embedded in OCT compound (Tissue-Tek), frozen in liquid nitrogen-cooled isopentane, and sectioned into 10 μm with CryoStar NX50 cryostat (Thermo Fisher scientific). Tissues sections were permeabilized with 0.3% (v/v) Triton-X 100 in 1X PBS for 20 min and blocked with 1% (w/v) BSA, 0.3 M glycine (Sigma, G7126), and 5% (v/v) goat serum (Cell signaling, 5425) in 1X PBS for 1 hour at room temperature. Tissue sections were then incubated with primary antibodies in blocking buffer at 4 °C overnight. After three 5 min washes with 1X PBS, secondary antibodies in blocking buffer, including Alexa Fluor Plus 488, 555, and 647 (Invitrogen), were pipetted to the tissue section for a 1 hour incubation at room temperature. Tissue sections were washed with 1X PBS thrice and incubated with DAPI for 20 min at room temperature. Tissue sections were then washed in 1X PBS for 5 min at room temperature and mounted with ProLong Gold antifade mountant (Invitrogen, P36930). Primary antibodies are listed in table S1.

### Single molecule fluorescence in *situ* hybridization (smFISH)

smFISH Probes were designed using NCBI primer-blast with 19–21 bp primer pairs for an amplicon length of 38–42 bp, primer melting temperature between 57 °C and 63 °C, and primer GC content between 35% and 70%. For each gene target, 7–13 sets of reverse complemented forward primers and reverse primers concatenated to flanking sequencing for HCR were ordered from IDT, mixed together, and diluted in nuclease-free water to create a split probe pool stock solution at 10 µM. Sequences of smFISH Probes are listed in table S2.

Collected tissues were immediately embedded in OCT compound, frozen in liquid nitrogen-cooled isopentane, and cryosectioned. Tissue sections were fixed in 4% formaldehyde for 12 min at room temperature, followed by 5 min wash in 1X PBS twice. Tissue sections were then incubated in 70% ethanol at room temperature for 1 hour to permeabilize the sections. Following 5 min wash in 1X PBS, 1 μM of smFISH probes in hybridization buffer constituted of 2X SSC, 5X of Denhart Solution, 10% Ethylene Carbonate, 10% Dextran Sulfate, and 0.01% SDS were then pipetted to the tissue section within a 9 mm X 9 mm Frame Seal chambers (Bio-rad, SLF0601) for incubation overnight at 37 °C inside a humidifying chamber. Tissue sections were washed in hybridization wash buffer constituted of 0.215 M NaCl, 0.02 M Tris HCl (pH 7.5), and 0.005 M EDTA for 20 min at 48 °C after removing the frame seal. Hairpin pairs labeled with Alexa Fluor 488, 546, and 647 (Molecular Instruments) were mixed, incubated at 95 °C for 1.5 min, and chilled for 30 min at room temperature before use. 0.06 μM fluorophore-labeled hairpins in amplification buffer constituted of 2X SSC, 5X of Denhart Solution, 10% Dextran Sulfate, and 0.01% SDS were pipetted to the tissue sections within a frame seal and incubated for 6 hours at room temperature. Tissue sections were washed in wash buffer for 20 min at 48 °C and incubated with DAPI for 20 min at room temperature. Tissue sections were then washed in 1X PBS for 5 min at room temperature and mounted with ProLong Gold antifade mountant.

### Calcium imaging

Cal520 (AAT Bioquest, 21130) was reconstituted in DMSO to prepare a 2 mM stock solution. NFVTs were removed from the PDMS post construct, mounted on silicone pillars (Electron Microscopy Science, 70338-40) glued over a confocal dish, and washed in M199 media without L-glutamine for 5 min twice. NFVTs were then incubated with 4 μM CAL520 in 0.04% w/v Pluronic F-127 (Sigma, P2443) and M199 media without L-glutamine for 90 min at 37 °C. Following two 5 min washes in M199 media without L-glutamine, NFVTs were equilibrated in M199 with HEPES modification and without L-glutamine (Sigma, M7528) for 20 min at room temperature.

Calcium signal of NFVTs was recorded using a Zeiss LSM 710 laser scanning microscope with a 10× water immersion objective under a resolution of 128 x 128 pixels and a zoom factor of 5 at 25 fps for 500 cycles. A custom MATLAB program was used to quantify the characteristics of calcium spikes.

### Ventricular engineered heart tissue imaging and analysis

Confocal images were acquired using a Zeiss LSM 710 laser scanning microscope with a 40× water immersion objective to obtain z-stacks of the tissue sections. ImageJ was utilized to analyze the z-stack images. Tissue coherency was measured using the ImageJ OrientationJ plugin. Sarcomere length was analyzed using software tool, SotaTool.

### Statistical analysis

Data was represented as means ± SEM. Statistical analysis was performed using Prism software (GraphPad). Significant differences between groups were analyzed by analysis of variance (ANOVA) with Bartlett’s multiple comparison test. Differences at P < 0.05 were considered statistically significant.

## Acknowledgments

This work was performed in part at the Cornell NanoScale Facility, a member of the National Nanotechnology Coordinated Infrastructure (NNCI), which is supported by the National Science Foundation (Grant NNCI-2025233), and Biotechnology Resource Center (BRC) Imaging Facility (RRID:SCR_021741) at the Cornell Institute of Biotechnology.

## Author contributions

Conceptualization: MLSP, GJS, JTB

Methodology: MLSP, GJS, JTB

Investigation: MLSP, AL, AD

Visualization: MLSP, GJS

Supervision: JTB

Writing—original draft: MLSP, JTB

Writing—review & editing: MLSP, JTB

## Data and materials availability

All data needed to evaluate the conclusions in the paper are present in the paper and/or the Supplementary Materials.

