## Supplemental data for "Finite afterload during asymmetrically cycled preload promotes fetal ventricular growth, maturation, and contractile function while suppressing fibrotic remodeling"

Science Advances

AAAS

Mong Lung Steve Poon *et al.*

**This PDF file includes:**

Figs. S1 to S6
Tables S1 to S2

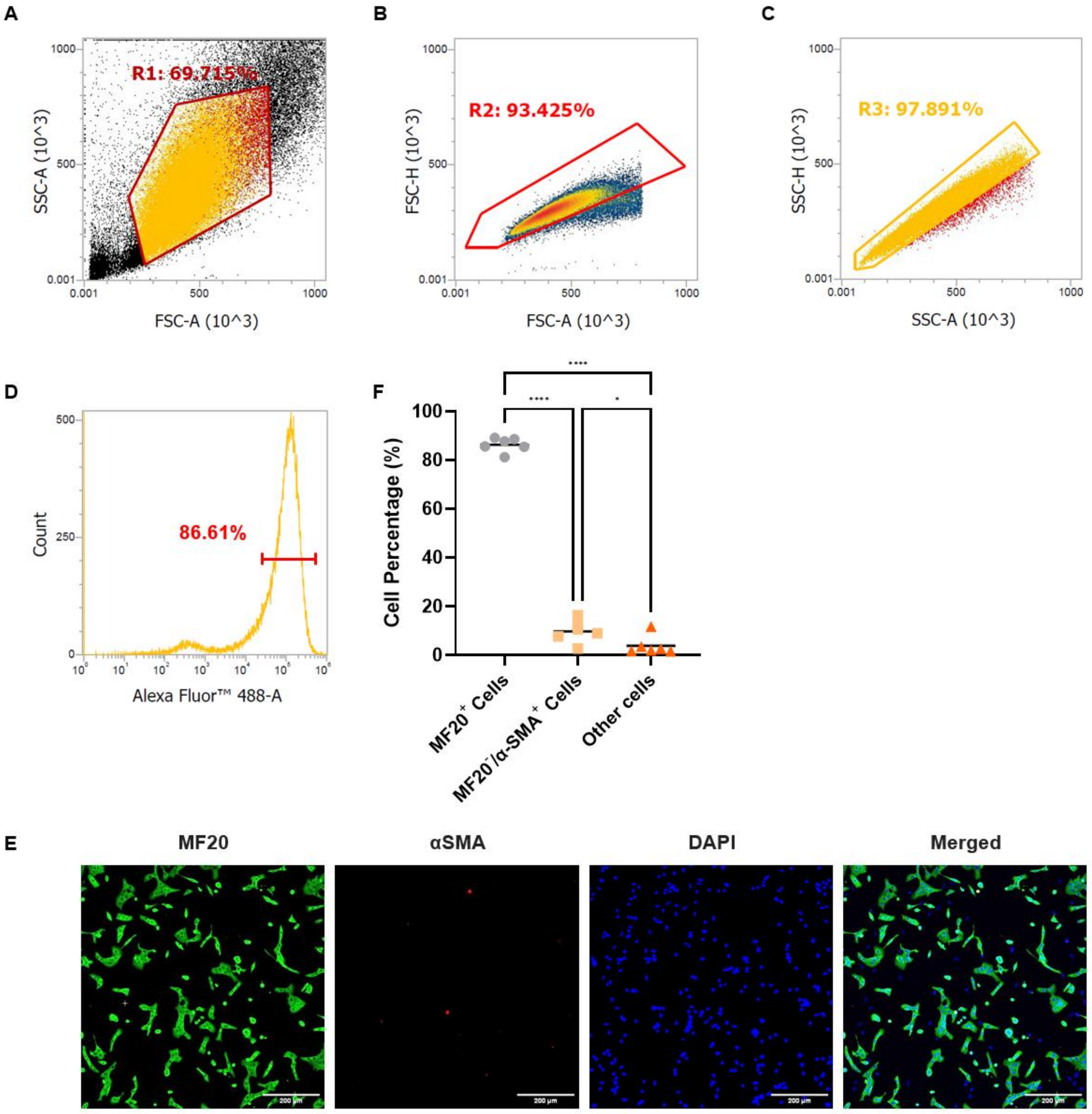


**Fig. S1. Fetal ventricular cells isolated from D7 chick embryo mainly composed of cardiomyocytes.**

(A-C) Cell gating and sorting of MF20 labelled chick heart ventricular cells following flow cytometry to ensure single cell. (D) Representative flow cytometry histogram of the sorted chick heart ventricular cells showing an enrichment of MF20+ cardiomyocyte at 86.61%. (E) Immunofluorescent images of chick heart ventricular cells seeded onto a cell culture dish for 1 day after isolation and purification and stained with MF20 (green), α-SMA (red), and DAPI (blue). Scale bar, 200 μm. (F) Cell percentage of chick heart ventricular cells. Data are means ± SEM, N = 3. One-way ANOVA with Tukey's test; $*P < 0.05$, $**P < 0.01$, $***P < 0.001$, and $****P < 0.0001$.

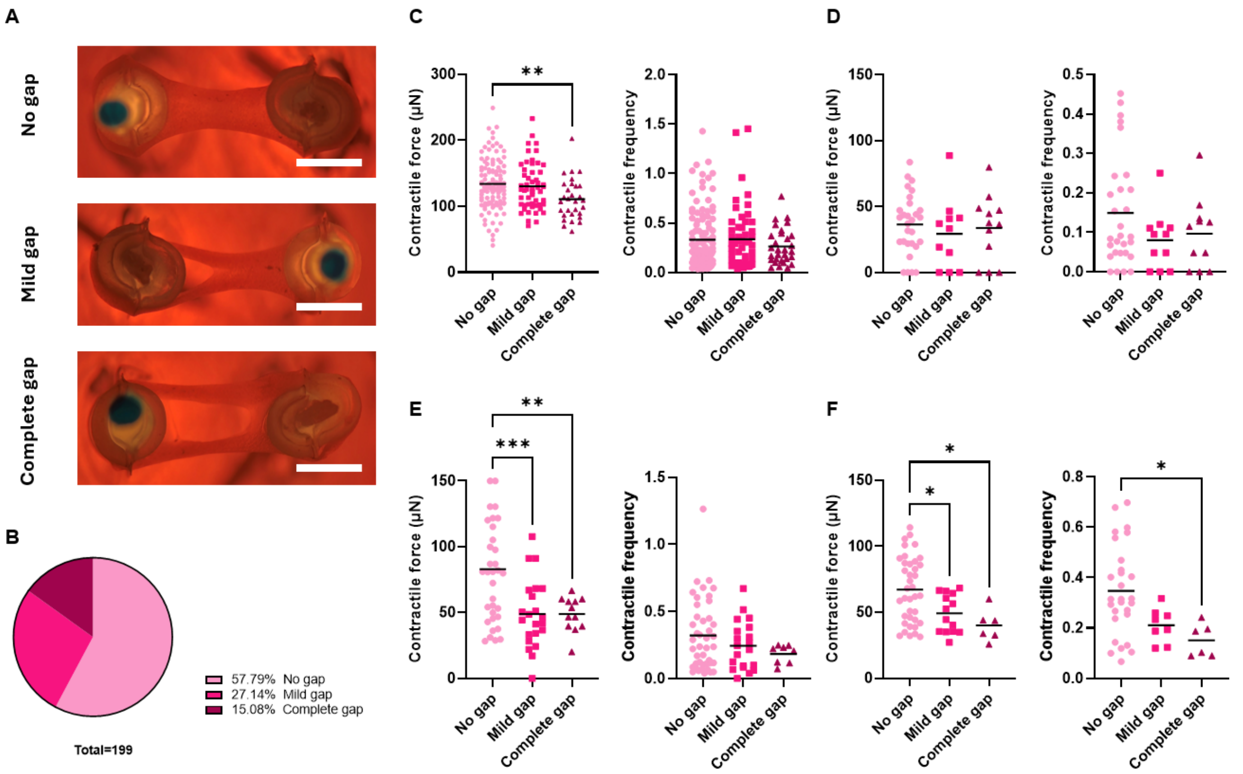


**Fig. S2. Gap formation in NFVT disrupts tissue contractility.**

(A) Representative microscopy images of NFVTs with no gap, mild gap, and complete gap formed. Scale bar, 2mm. (B) Pie chart of the occurrence of NFVTs with no gap, mild gap, and complete gap. Contractile force and contractile frequency of NFVTs with no gap, mild gap, and complete gap on (C) Day 5 and on Day 8 of (D) SC, (E) HA, and (F) LA mechanical stimulation regimens. All data are means ± SEM, N = 6. One-way ANOVA with Tukey's est; *P < 0.05, **P < 0.01, ***P < 0.001, and ****P < 0.0001.

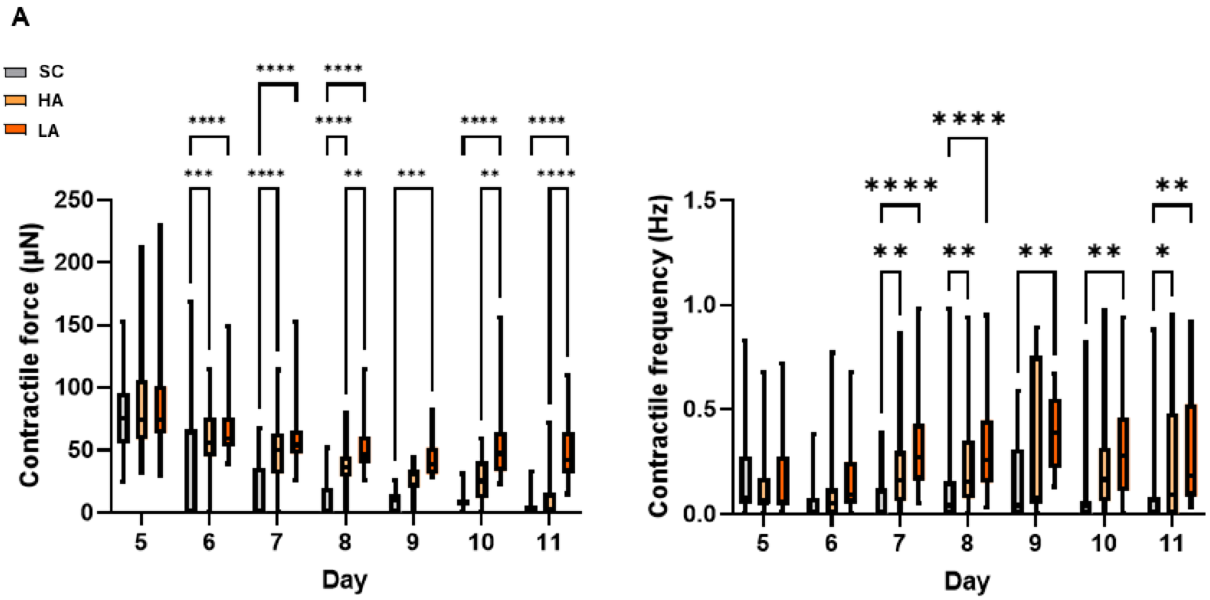


**Fig S3. Contractile force and frequency of NFVTs across the stimulation period.**

(A) Normalized contractile force and (B) normalized contractile frequency of NFVTs relative to SC from Day 5 to Day 11 under SC, HA/1Hz, LA/1Hz and LA/2Hz mechanical stimulation regimens. Data are means ± SEM, N = 6. Two-way ANOVA with Tukey's test; *P < 0.05, **P < 0.01, ***P < 0.001, and ****P < 0.0001.

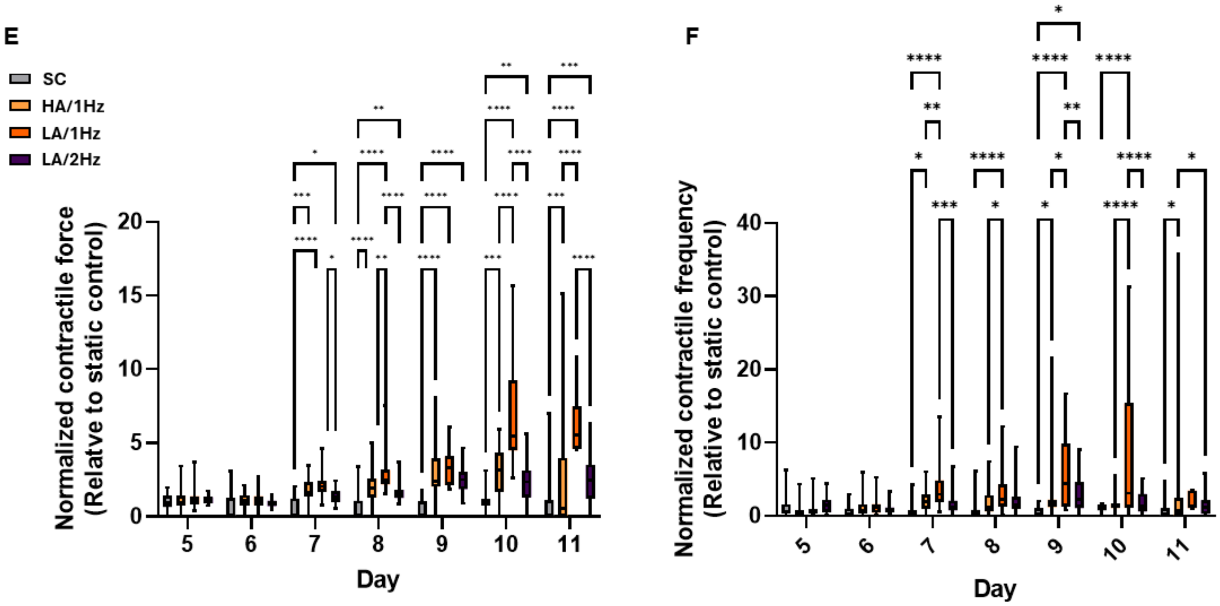


**Fig S4. LA at 2 Hz suppresses the improvement of tissue contractility**

(A) Normalized contractile force and (B) normalized contractile frequency of NFVTs relative to SC from Day 5 to Day 11 under SC, HA/1Hz, LA/1Hz and LA/2Hz mechanical stimulation regimens. Data are means ± SEM, N = 6. Two-way ANOVA with Tukey's test; *P < 0.05, **P < 0.01, ***P < 0.001, and ****P < 0.0001.

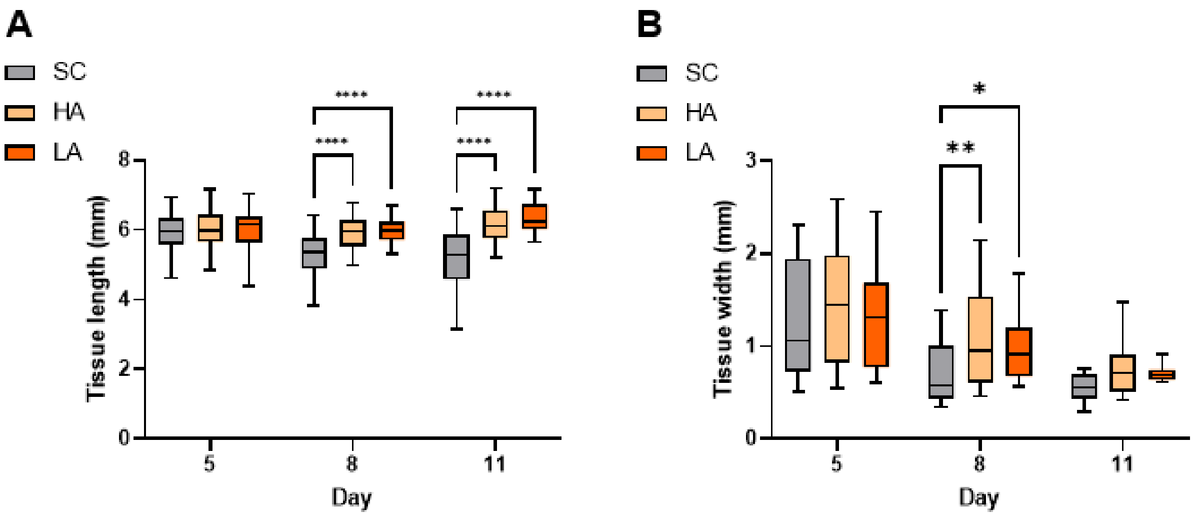


**Fig S5. LA upheld tissue length and tissue width**

(A) Tissue length and (B) tissue width of NFVTs on Day 5, 8, and 11 of SC, HA, or LA mechanical stimulation regimens. Data are means ± SEM, N = 6. Two-way ANOVA with Tukey's test; *P < 0.05, **P < 0.01, ***P < 0.001, and ***P < 0.0001.

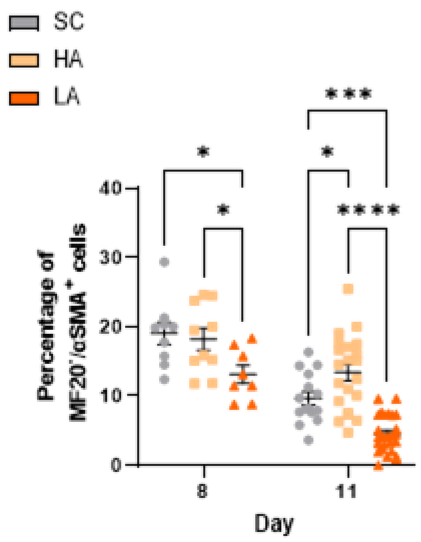


**Fig S6. Fig. S6. HA led to higher cardiac fibroblast abundance on Day 11.**

Percentage of cardiac fibroblast within NFVTs on Day 8 and 11 of SC, HA, or LA mechanical stimulation regimens. Data are means ± SEM, N = 6. Two-way ANOVA with Tukey's test; *P < 0.05, **P < 0.01, ***P < 0.001, and ***P < 0.0001.

| **Table S1: Immunolabeling** | | | |
|---|---|---|---|
| **Protein** | **Supplier** | **Clone/ Cat no.** | **Clonality** |
| Myosin 4 (MF20) | Thermo Fisher Scientific | 14-6503-82 | moAb |
| Alpha smooth muscle actin | Proteintech | 55135-1-AP | poAB |
| Sarcomeric Alpha Actinin | Abcam | ab9465 | moAb |
| Connexin 43 (GJA1) | Abcam | ab11370 | poAB |
| PIEZO1 | Cell signaling Technology | #62649 | moAb |
| YAP1 | DSHB | AB_2619554 | moAb |
| Activated Notch1 | Abcam | ab52301 | poAB |
| Cleaved Caspase-3 (Asp175) | Cell signaling Technology | #9661 | poAB |
| Phospho-Histone H3 (Ser10) | Cell signaling Technology | #9701 | poAB |

Table S2: smFISH probes

| Gene symbol | Probe pair number | Initiator 1 | Target Split Probe 1 | Target Split Probe 2 | Initiator 2 |
|---|---|---|---|---|---|
| TNNT1 | 1 | gTCCCTgCCTCTATATCTTT | CAACGACCTCTTCAGAGTCCG | TGCTCCTGCTCGTATTCCTC | TTCCACTCAACTTTAACCCg |
| TNNT2 | 2 | gTCCCTgCCTCTATATCTTT | GCATGAAGGGCTTGGGTTTG | ATTTTGGGAGGCACCAGGTT | TTCCACTCAACTTTAACCCg |
| TNNT2 | 3 | gTCCCTgCCTCTATATCTTT | AAATGGGCTTCGATGAGGGC | TCCTCCTTCTTCCTGCTCTCA | TTCCACTCAACTTTAACCCg |
| TNNT2 | 4 | gTCCCTgCCTCTATATCTTT | GCATGTAGCCTCCAAAGTGC | CCACCCTTCTTCTCCGACTT | TTCCACTCAACTTTAACCCg |
| TNNT2 | 5 | gTCCCTgCCTCTATATCTTT | TTCTTCTCCCGCTCCGTTTG | CGCTCGCTGAGGATCTTTTT | TTCCACTCAACTTTAACCCg |
| TNNT2 | 6 | gTCCCTgCCTCTATATCTTT | GTGGTCGATGTTCAGAGGCT | CCCTCAGTTTGTCTTCGCTGA | TTCCACTCAACTTTAACCCg |
| TNNT2 | 7 | gTCCCTgCCTCTATATCTTT | GATGGTTTGCCACAGCTCCT | ATTTCTCAGCCTCCAGGTCAC | TTCCACTCAACTTTAACCCg |
| TNNT2 | 8 | gTCCCTgCCTCTATATCTTT | TCTCGTACTTCTGCCGCTTG | ACACGATTTCGAAGGACGTTG | TTCCACTCAACTTTAACCCg |
| TNNT2 | 9 | gTCCCTgCCTCTATATCTTT | CCTTCTGGTGGTCACTGAC | GCAGCCTTTGACCCTTTGAC | TTCCACTCAACTTTAACCCg |
| TNNT2 | 10 | gTCCCTgCCTCTATATCTTT | TCAAGCGGCAGAAGTACGAG | ACGATTTCGAAGGACGTTGA | TTCCACTCAACTTTAACCCg |
| FN1 | 1 | CCTCgTAAATCCTCATCAAA | CCAACGTGTTCCCCAGGTAA | TCCACCGTAGCAGGTACAGA | AAATCATCCAgTAAACCgCC |
| FN1 | 2 | CCTCgTAAATCCTCATCAAA | TTCACCTCTGCCGTTACCAG | GTGTGCCTTTCGCATTTCCA | AAATCATCCAgTAAACCgCC |
| FN1 | 3 | CCTCgTAAATCCTCATCAAA | AAGGCAGAACACAGGCTTCT | AGTCCTCCCGTTGTAGGTGA | AAATCATCCAgTAAACCgCC |
| FN1 | 4 | CCTCgTAAATCCTCATCAAA | ACTGATCAACGGGATCGCAC | ACGAGTTTCGGAATCCTGGC | AAATCATCCAgTAAACCgCC |
| FN1 | 5 | CCTCgTAAATCCTCATCAAA | CCTTCCAGTGCCTTCCAGAG | GTGACCAGGGATGGTAGCTT | AAATCATCCAgTAAACCgCC |
| FN1 | 6 | CCTCgTAAATCCTCATCAAA | GGGGTCTGCTCCAGCTAATG | GTAGCCTGTGATTGGAGCCT | AAATCATCCAgTAAACCgCC |
| FN1 | 7 | CCTCgTAAATCCTCATCAAA | CTTCCACCCGATAGCCAGAC | AGGGCGATTGACAGGGATTG | AAATCATCCAgTAAACCgCC |
| FN1 | 8 | CCTCgTAAATCCTCATCAAA | GTATGTCCGTGGCTGACCTC | GTAGCAGAAGGGCCCACATT | AAATCATCCAgTAAACCgCC |
| FN1 | 9 | CCTCgTAAATCCTCATCAAA | CCAAGAAACTGCTCAGCACG | CTTCACAGGTGAGTAGCGCA | AAATCATCCAgTAAACCgCC |
| FN1 | 10 | CCTCgTAAATCCTCATCAAA | TGTCACCCGATACCCTGTGA | TGTCCTTTCTTGGGGACAGC | AAATCATCCAgTAAACCgCC |
| FN1 | 11 | CCTCgTAAATCCTCATCAAA | TCCATCCTCAGGGCTTGAGT | GCCGGCAATAGCTCATGGAT | AAATCATCCAgTAAACCgCC |
| FN1 | 12 | CCTCgTAAATCCTCATCAAA | ATGTGCTGTCCGGAGAAAGG | TAATCCCGACACGACAGCAG | AAATCATCCAgTAAACCgCC |
| FN1 | 13 | CCTCgTAAATCCTCATCAAA | TATTGTCTGTGGGTGCCCTG | ACTTGCTGACCTTGGTGTCC | AAATCATCCAgTAAACCgCC |
| FN1 | 14 | CCTCgTAAATCCTCATCAAA | GAAGTGCCAGGAACCCTGAA | CAGTGAGCGTAGCACTGGAG | AAATCATCCAgTAAACCgCC |
| TGFB2 | 1 | gAggAgggCAgCAAACggAA | ACGCTCAGGAGATAGCAGTG | CCAGATCCAGGGTGAGGAAC | TAgAAgAgTCTTCCTTTACg |
| TGFB2 | 2 | gAggAgggCAgCAAACggAA | GCAGGTAGACAGGCTGAGAG | TGATCCATGTCGAGGGTGCT | TAgAAgAgTCTTCCTTTACg |
| TGFB2 | 3 | gAggAgggCAgCAAACggAA | GATGGAGATGACCTCCGGGG | AGGTCCCTGGTGCTGTTGTA | TAgAAgAgTCTTCCTTTACg |
| TGFB2 | 4 | gAggAgggCAgCAAACggAA | GTGGTTGGCTTTCTCCTGCA | CTCTCGCAAGTGGCAGCTCT | TAgAAgAgTCTTCCTTTACg |
| TGFB2 | 5 | gAggAgggCAgCAAACggAA | TCGGGGTAAAAAGGCTGCAT | AGCTTGGTGGGATGGCATTT | TAgAAgAgTCTTCCTTTACg |
| TGFB2 | 6 | gAggAgggCAgCAAACggAA | GGAAGACCCTGAACTCAGCC | CGCCTTTGAGTTCTGCAGGC | TAgAAgAgTCTTCCTTTACg |
| TGFB2 | 7 | gAggAgggCAgCAAACggAA | AGCCATTCATGTACAGCCTC | GGTTCCTGTCTCTGTGATGG | TAgAAgAgTCTTCCTTTACg |
| TGFB2 | 8 | gAggAgggCAgCAAACggAA | GACTGGGCTGTTGCGACTCA | TAGAGCACGCTTCTTCCGCC | TAgAAgAgTCTTCCTTTACg |
| TGFB2 | 9 | gAggAgggCAgCAAACggAA | AAATCCTGGGACACGCAGCA | AGAGGATGGTGAGGGGCTCT | TAgAAgAgTCTTCCTTTACg |

| Gene symbol | Probe pair number | Initiator 1 | Target Split Probe 1 | Target Split Probe 2 | Initiator 2 |
|---|---|---|---|---|---|
| POSTN | 1 | gTCCCTgCCTCTATATCTTT | GTCGCGTGCCCTTATTCGAC | GCACAGACATTTGGGCCTTG | TTCCACTCAACTTTAACCCg |
| POSTN | 2 | gTCCCTgCCTCTATATCTTT | TCCCACAATACCAAGAGTGCC | TACTGCTGAGTGGAGGTAGC | TTCCACTCAACTTTAACCCg |
| POSTN | 3 | gTCCCTgCCTCTATATCTTT | TGATCGCTTCAGAGCACTGA | TTCATAGACAGCTCCACCCA | TTCCACTCAACTTTAACCCg |
| POSTN | 4 | gTCCCTgCCTCTATATCTTT | CTTGCTTGGCAGAATCAGGAA | GCACCTCCAAGCTCAATGAC | TTCCACTCAACTTTAACCCg |
| POSTN | 5 | gTCCCTgCCTCTATATCTTT | ACCAGGAGAGTGTATTGGCC | TGAGAAAGCACGATTCTGTGG | TTCCACTCAACTTTAACCCg |
| POSTN | 6 | gTCCCTgCCTCTATATCTTT | AGGCCGTGACAGAACATCAT | ACAAACAGAGTCCATGCTCCA | TTCCACTCAACTTTAACCCg |
| POSTN | 7 | gTCCCTgCCTCTATATCTTT | AACTCCTTGTGTCAGGTGGT | GCTCAAAGCCACTTCCAATGA | TTCCACTCAACTTTAACCCg |
| POSTN | 8 | gTCCCTgCCTCTATATCTTT | CTGATCGTTTCCAACAGGCA | GCTTCTTCAGGATTGTGAGCA | TTCCACTCAACTTTAACCCg |
| POSTN | 9 | gTCCCTgCCTCTATATCTTT | CGCCCATCAATGATTTTGGTG | CTCTGTCACTTCCACAGGTCG | TTCCACTCAACTTTAACCCg |
| POSTN | 10 | gTCCCTgCCTCTATATCTTT | GCTCCCTGCAGAAGTCTTTTG | ACCTTGGTGTACTCAGTTCCA | TTCCACTCAACTTTAACCCg |
| POSTN | 11 | gTCCCTgCCTCTATATCTTT | CCCTTGTGTCCTCTTGTTTGC | GTCCTCCTTCTTGCTGATCCC | TTCCACTCAACTTTAACCCg |
| COL5A2 | 1 | gAggAgggCAgCAAACggAA | AGACCTTGTGATCCAGCTGG | CTGTCCCTTCCGTACCAGGA | TAgAAgAgTCTTCCTTTACg |
| COL5A2 | 2 | gAggAgggCAgCAAACggAA | TTCACCTATGGGGCCAATGG | CGAGGGCCTCTTTTCCCTTC | TAgAAgAgTCTTCCTTTACg |
| COL5A2 | 3 | gAggAgggCAgCAAACggAA | GAGCACCCCTCTCTCCAACT | TGGAAAACCACGGTTCCCAG | TAgAAgAgTCTTCCTTTACg |
| COL5A2 | 4 | gAggAgggCAgCAAACggAA | CTATCGAACCTGCAGGACCA | GCCTGGCTGACCTCTGATTC | TAgAAgAgTCTTCCTTTACg |
| COL5A2 | 5 | gAggAgggCAgCAAACggAA | TTGTCCTGGAACACCTGCAT | AAGCCAGTCACACTCACTCT | TAgAAgAgTCTTCCTTTACg |
| COL5A2 | 6 | gAggAgggCAgCAAACggAA | TCACCTGCAAGTCCTGATGC | GACCTCCAGCCATACCCTTT | TAgAAgAgTCTTCCTTTACg |
| COL5A2 | 7 | gAggAgggCAgCAAACggAA | ACCTCTTGCCCCATCATTGC | GGGCCTATTGGACCAGGAAG | TAgAAgAgTCTTCCTTTACg |
| COL5A2 | 8 | gAggAgggCAgCAAACggAA | CCCTGTCACCTCGATCTCCA | ACCTCTGTGTCCTTTCTGGC | TAgAAgAgTCTTCCTTTACg |
| COL5A2 | 9 | gAggAgggCAgCAAACggAA | CAAGTCCGTGCTGGGTGTTT | GGCACAGCTTCAGGTCATCA | TAgAAgAgTCTTCCTTTACg |
| COL5A2 | 10 | gAggAgggCAgCAAACggAA | AGGTCATTGGCCCCTTTCAG | TTCCCTCCGCCTTGATCTCC | TAgAAgAgTCTTCCTTTACg |
| COL5A2 | 11 | gAggAgggCAgCAAACggAA | CGTTCTGCGTTCGGTACTCA | TATGATAGGCAACCGTGCCA | TAgAAgAgTCTTCCTTTACg |
